# Polygenic architecture of adaptation to a high-altitude environment for *Drosophila melanogaster* wing shape and size

**DOI:** 10.1101/2024.02.08.579525

**Authors:** Katie Pelletier, Megan Bilodeau, Isabella Pellizzari-Delano, M. Daniel Siemon, Yuheng Huang, John E. Pool, Ian Dworkin

## Abstract

As is typical of small insects, populations of *Drosophila melanogaster* adapted to high altitude environments evolved increased body size, disproportionality large wings, and differing wing shape compared to low-altitude ancestors. In one instance the colonization of high-altitude environments in Ethiopia is recent (2000-3000 years ago), and is a useful system to study alleles contributing to adaptive divergence. Unlike predictions derived from formulations Fisher-Kimura-Orr geometric model based on *de novo* mutations concurrent with selection, recent models predict segregating alleles in a population are more likely to contribute to adaptation on short time scales, particularly when populations are large and genetically diverse, like *D. melanogaster*. Strains derived from lowland (∼500m above sea level – ASL) and highland (∼3000m ASL) populations were used to generate F20 advanced-intercrosses. From each cross, phenotypically extreme individuals for size and shape were pool-sequenced, and genetic differentiation among pools of individuals demonstrated a polygenic architecture of divergence for size and shape. We identified one QTL of large effect, contributing to adaptive divergence in shape. This QTL is not observed in all crosses, pointing to the importance of examining independent genetic backgrounds when mapping alleles contributing to adaptation. Despite the intrinsic links between shape and size, we find a unique genetic basis of adaptation for these traits. This work demonstrates that many alleles, throughout the genome, rather than single, large effect alleles, contribute to adaption for *Drosophila* wing shape and size, adding to the growing body of evidence for polygenic adaptation.

## Introduction

Identifying the genetic basis of heritable variation contributing to adaptive divergence is an important first step in evolutionary genetics. This creates the opportunity for future studies, whether examining the mechanisms of allelic effects on development or metabolism, allelic dynamics over time, or inheritance patterns, among other questions that can be studied. However, for many of the traits of interest to evolutionary biologists, mapping the alleles contributing to divergence is not a case of simply identifying a single segregating locus of large phenotypic effect. For adaptively divergent traits, polygenicity of complex traits is common. Even traits where alleles of large phenotypic effect are identified, there are often also many genes with alleles of small effects also involved. Several studies have mapped alleles contributing to trait divergence due to adaptation and have identified one or a small number of major effect alleles contributing to phenotypic divergence. For instance, alleles at *mc1r* influencing coat colour in mice accounting for ∼36% of the trait variation (Hoekstra, 2006; Hubbard et al., 2010), or *Eda* influencing stickleback armor plate variation (Barrett et al., 2008; Colosimo et al., 2004; Hubbard et al., 2010). Such variants of large effect are often a result of *de novo* mutations that occurred concurrently with selection, or, were very rare in ancestral populations (Dittmar et al., 2016). It is important to recognize that there is a form of “selection bias” at play with these studies, which can in turn bias our understanding of the genetic architecture of adaptation. In those systems where allelic variations of large phenotypic effect are present, the ability to map those variants is comparatively straightforward, requiring relatively modest sample sizes. These examples may represent a very biased sample of the genetic architecture of complex traits, as identifying these single genes of large effect requires not only smaller sample sizes but also smaller numbers of unique genotypes to identify alleles of large effect. In such studies, particularly those with smaller mapping populations, alleles of modest phenotypic effect are likely to be overlooked or remain unidentified. With the falling cost of genomic sequencing and more widespread whole genome mapping in larger populations, it has become apparent that adaptive changes in trait values generally involve contributions from many loci (Pritchard and Di Rienzo, 2010). The small effect variants across the loci usually include a substantial fraction of alleles segregating in ancestral populations at non-negligible frequency, as opposed to *de novo* mutations, or very rare variants. The relative contribution of large and small effect alleles matters, as they will affect relative contribution of (hard) genetic sweeps at a few genomic loci at one end of the spectrum to subtle shifts in allele frequencies across many loci on the other. The former case is much easier to detect in population genetics data than the latter, as hard sweeps leave clear genomic footprints of low nucleotide diversity. As such, where on the spectrum of the effect size distribution the alleles contributing to adaptation occur, and how this depends on features of the ancestral population and the relevant traits (mutational target size and segregating variation for the trait, *N_e_*, etc), is an open but essential question in understanding the mechanisms of adaptation for complex traits (Barghi et al., 2020).

Hard sweeps from single alleles of large effect, subtle shifts in allele frequencies and all the points on the spectrum allow for rapid adaptation with contribution from multiple loci, but the relative effect sizes of contributing alleles and contribution of mutation and segregating variation varies among them. The theory of an adaptive walk based on the extensions of the Fisher-Kimura geometric model predicts subsequent fixation of alleles (Fisher, 1930; Kimura, 1968; Orr, 2005, 1998). This sweep can be “hard”, due either to *de novo* mutations contributing (initial frequency of 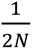), or selection “capturing” a single haplotype with a rare variant. Alternatively the sweep can be “soft”, with contributions from alleles segregating (>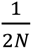) in the population that are already present on multiple, distinct haplotypes (Coop and Ralph, 2012). For alleles to contribute to adaptation of the population, they must be sufficiently beneficial (or high enough frequency for segregating variants) to overcome stochastic effects of genetic drift. Populations with large, long-term effective population size (*N_e_*) tend to have high genetic diversity, and often have ample standing genetic variation for a trait that ultimately contributes to adaptive evolution. In such cases, the rate of allele frequency change at individual variants of large effect will be reduced in favor of polygenic adaptation due to modest shifts in allele frequency across many loci (Chevin and Hospital, 2008). As predicted by Fisher’s infinitesimal model and subsequent extensions, the predicted effect of most mutations is small because of high pleiotropy resulting in most large effect alleles being too deleterious to segregate in populations (Boyle et al., 2017; Fisher, 1930). Small shifts in allele frequency across many loci in response to directional selection, followed by a period of stabilizing selection, can result in phenotypic divergence, without fixation of alleles among diverged populations (Hayward and Sella, 2019). Because of the small shifts predicted by this model, identification of particular alleles will be difficult, as approaches like *F_ST_* scans to identify differentiated alleles may not be able to detect subtle shifts (Yeaman, 2015). Study design in regard to diversity of founding strains and size of mapping populations can have an effect of the success of identification of alleles and may even bias inferences made about polygenicity of a trait. Approaches using experimental mapping populations with multiple strains derived from relevant natural populations may be better suited to identify alleles contributing to adaptive divergence, as small effect alleles will be more likely to be identified and polygenicity inferred.

*Drosophila melanogaster* populations are a particularly tractable system for mapping alleles contributing to phenotypic divergence, due to: short generation times, ease of culture and the extensive genetic tools available. It is an equally useful system, as its world-wide distribution has resulted in numerous cases of both seasonal (Machado et al., 2021; Yu and Bergland, 2022) and local adaptation, with respect to latitudinal (Bergland et al., 2016; de Jong and Bochdanovits, 2003; Land et al., 1999) and altitudinal (Bastide et al., 2014; Fabian et al., 2015; Pitchers et al., 2013; Pool et al., 2016) variation. As is typical of small insects adapting to life at high altitude (Dillon, 2006; Dillon and Frazier, 2006), numerous traits vary between high and low altitude populations of *D. melanogaster* in sub-Saharan Africa, including body size, pigmentation, wing shape and size (Bastide et al., 2016; Fabian et al., 2015; Groth et al., 2018; Lack et al., 2016, 2016; Pitchers et al., 2013; Pool, 2016). During adaptation to high altitude environments, small insects typically show a suite of evolved changes such as increased pigmentation, body size, and disproportionate increases in wing size, as wing loading is a key variable in the ability to fly in the thin and cool air of high altitude environments (Dillon, 2006). Flight performance can evolve rapidly in *D. melanogaster* (Marden et al., 1997). High altitude *D. melanogaster* have a larger wing size and increased wing loading, compared to lowland populations (Fabian et al., 2015; Lack et al., 2016; Pitchers et al., 2013). Using nearly isogenic strains isolated from a highland population (Fiche, Ethiopia >3000 m ASL) and a lowland population representing the putative ancestral range (Siavonga, Zambia 500 m ASL) (Lack et al., 2015; Sprengelmeyer et al., 2020), mapping studies have been conducted to identify the genetic basis of traits contributing to adaptive divergence. In the case of pigmentation, a small number of QTN have been identified accounting for the majority of pigmentation variation between populations, with a number of alleles of smaller effect contributing (Bastide et al., 2014). In contrast, mapping studies that have examined differences in body and wing size identified a polygenic basis with little evidence of selective sweeps at single loci and no single QTNs identified (Sprengelmeyer et al., 2022). The differences in the genetic architecture of these traits could help to explain the differences in the two patterns discovered. Pigmentation has a ‘simpler’ genetic basis, as a result of a smaller mutational target size (Dembeck et al., 2015; Wittkopp et al., 2003) than wing size, where more than 15% of genes (Carreira et al., 2013, 2009; Houle and Fierst, 2013) can contribute to the mutational variance. The previous size mapping study used one Zambian and four Ethiopian derived strains to identify alleles, this can create a bias in the results as only one genome from the Zambian population is used to represent this population. By adding mapping studies with capturing genetic diversity within each divergent population, while also improving mapping power with larger numbers of phenotyped individuals per cross, we can begin to understand the degree of polygenicity for adaptation of wing size and size.

In contrast with size, the genetic basis of divergence for wing shape between these two populations has not been investigated, although there are well-documented wing shape differences, even after accounting for allometry, between highland and lowland populations (Pesevski and Dworkin, 2021; Pitchers et al., 2013). Like wing size, wing shape has a large mutational target size (Carreira et al., 2011; Houle et al., 2017; Weber et al., 2005, 2001) but differs from wing size in its multidimensional nature. Wing shape can vary in many directions; as the number of available dimensions of variation increases, the likelihood of genetic effects being aligned by chance decreases. Both simulations and empirical studies have demonstrated that taking an explicitly multivariate approach to multidimensional traits (rather than trying to define them as a series of principal components, for instance) is considerably more powerful for understanding the genetic architecture of complex traits (Pitchers et al., 2019; Porter and O’Reilly, 2017). The increased number of directions in which effects can vary allows for the analysis of shared directions of variation. The major axes of wing shape variation between Drosophild species are relatively well aligned with mutational variation within *D. melanogaster* (Houle et al., 2017). This example is on a much longer time scale than the relatively recent adaption to high altitude observed between highland and lowland populations (<3000 years) (Sprengelmeyer et al., 2020) but provides a motivating model with which to understand the genetic architecture of divergence on short and long time scales.

Allometry is a major component of shape variation in many systems, including *Drosophila* wings (Gidaszewski et al., 2009; Klingenberg, 2016; Sztepanacz and Houle, 2021). Although it is often possible to partition allometric and non-allometric components of shape (Klingenberg, 2022), and there is evidence for a change in the non-allometric component of shape between highland and lowland populations (Pitchers et al., 2013), it is not clear if the change is a consequence of selection on shape itself or due to selection for larger wings though allometry. The adaptive benefit of larger wings at high altitudes is, at least partly, explained by the increased surface area of the wing creating lift at lower air pressures as lower air pressures can inhibit flight performance of wild-caught flies (with smaller wing area compared to Ethiopian populations) in North America (Dillon, 2006; Dillon and Frazier, 2006). However, the adaptive benefits of differences in wing shape is not currently well understood, in part because it remains difficult to test (but see: Ray et al., 2016). Recent work suggests that variation in aspects of wing morphology beyond that of wing area can contribute to flight performance (Flaibani et al., 2023). By asking if variation for the non-allometric components of wing shape VS wing size have partially independent genetic architectures, we can test whether they are genetically separable.

Using strains derived from highland and lowland populations of *D. melanogaster*, we investigated the genetic architecture of two complex traits, wing shape and size. We demonstrated a polygenic architecture of divergence for wing size and wing shape, with few QTL shared between different crosses, supporting the hypothesis that alleles contributing to variation for these traits are not fixed in the derived, high-altitude population. Despite the partially independent genetic basis of adaptation among strains, a common allometric size-shape relationship and similar structure of shape variation was observed between different F20 intercrosses, even when the parental inbred lines the crosses were derived from have distinct allometric patterns. We also demonstrated the partially independent genetic basis of wing shape and size adaptation, despite the intrinsic link though the size-shape allometric relationship. Overall, this study provides support for the model of polygenic adaptation due to small shifts in allele frequency rather than subsequent fixations of alleles.

## Methods

### Generation of F20 advanced intercross populations

To map the genetic basis of shape and size adaptation between low altitude (Siavonga, Zambia, “ZI,” 16.54 S, 28.72 E, 530 m ASL) and high altitude (from Fiche, Ethiopia, “EF,” 9.81 N, 38.63 E, 3,070 m ASL) populations of *Drosophila*, 3 highland lines and 3 lowland lines were selected for generating pairwise, F20 advance intercrosses. The Zambian (ZI192N, ZI251N, ZI418N) and Ethiopian (EF43N, EF81N, EF96N) lines were selected from wild-caught isofemale lines that were inbred for 8 generations (Lack et al., 2015)(Figure 1). Lines were selected to be free of common, known chromosomal inversions*: In(1)A, In(1)Be, In(2L)t, In(2R)NS, In(3L)OK, In(3L)P, In(3R)K, In(3R)Mo*, and *In(3R)P* (Lack et al., 2016, 2015). All crosses were done at 20°C in the Pool lab (University of Wisconsin), using the lab’s standard recipe (see supplementary file 1 for all fly media recipes). Each pairwise cross was done in a single 28cm x 14cm x 15cm cage. In the first generation, 8 parents from the appropriate highland and lowland lines were crossed in a reciprocal manner to ensure equal contributions of X chromosomes from high and low-land populations. 125 F1 individuals of each sex from each reciprocal cross were selected to start the cage cross population. In cages, flies were provided with 14 vials of food and allowed to lay for 1 week before food was changed. For each non-overlapping generation, approximately 1,200 progeny from the previous generation produced the next generation. No selection was used in choosing which individuals contributed to the next generation. After 20 generations of crossing, adult flies were collected and stored in 70% ethanol and shipped to McMaster University for phenotyping, DNA extraction and sequencing.

**Figure 1.**
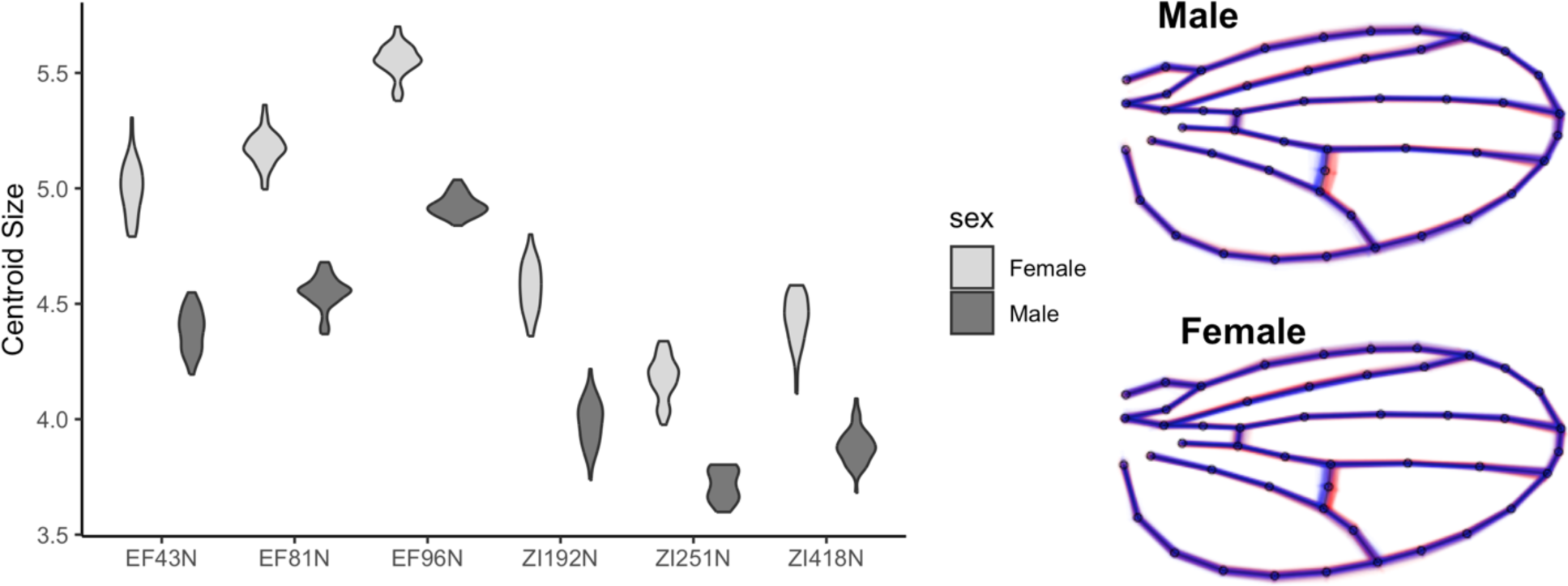
Divergence for wing size and shape, within and between high and low altitude strains used for mapping. (Left) Distribution of wing sizes, measured by the centroid size of the wing, for parental strains used for crosses. Ethiopian derived strains are on average larger than those from derived from Zambia. (Right) Distribution of individual wing shapes, within sex, between high (red) and low (blue) altitude populations. Approximately 150 wings are plotted for each sex and population.

### Collection of Morphometric Data

The right wings from individual flies were dissected and mounted in 70% glycerol in PBS, while bodies were stored individually in 95% ethanol in wells of a 96 well plate until genomic extraction. Wings were imaged using an Olympus DP30B camera mounted on an Olympus BX51 microscope (Olympus software version 3.1.1208) using a 4X objective lens (total 40X magnification) and images were taken using cellSens Standard (version 1.14).

Shape data were collected using two methods, with the second method used to increase the speed of data collection, and these methods largely captured a similar shape change (Figure S1). The first method (referred to as the WINGMACHINE method) used extracted positional information for 14 landmarks and 34 semi-landmarks, allowing for 58 available dimensions of variation (explained below). In this method, shape data were collected using the WINGMACHINE pipeline outlined in (Houle et al., 2003). First, two landmarks were placed at the humeral break and alula notch using tpsDIG2 (version 2.16). Wings (Van der Linde 2004–2014, v4) software was used to fit nine cubic B-splines, and to manually correct errors in spline positions. All shape data was subjected to Procrustes superimposition to scale landmark configurations to the same size, translate to the same location and registration (rotation) to minimize distances between landmark configurations (Rohlf and Slice, 1990). Landmark and semi-landmark data was extracted using the program CPR (version 1.11). While this produces 48 landmark pairs, translation (2), scaling (1) and registration (1) results in the loss of 4 dimensions of variation, while semi-landmark (34) variation is constrained to vary along (approximately) a single axis (dimension). This results in 24 + 34 = 58 dimensions of variation.

The second method (referred to as the 15-landmark method) extracted positional information for 15 landmarks (no semi-landmarks). 4 dimensions of variation are constrained due to translation, scaling and registration of landmark configuration resulting in 26 available dimensions of variation. Landmarks were identified using a Fiji (v4.0.3) plugin (courtesy of Dr. C. Klingenberg) designed to rapidly capture wing shape. In this case, Procrustes superimposition was done with the gpagen() function in geomorph (v 4.0.3), (Adams and Otárola-Castillo, 2013). The shape changes captured using both methods is similar, as demonstrated by the similar variance structures and shape changes between the two methods (Figure S1, S2). In both cases, the methods used for downstream analysis of shape were the same.

Males and females were considered separately for downstream analyses due to the considerable sexual dimorphism for wing shape and size. To partition allometric components of shape in the data, and to ensure that the size and shape axes of variation were independent, a PCA that included the shape residuals along with the natural log transform of centroid size was used. Using this method, most of the size variation, including the allometric component of shape, is captured by PC1 (Klingenberg, 2022) (Figure S3). The remaining PCs (PC2 – 58) for the WINGMACHINE method, or PC2 – PC26 for the 15-landmark method were used for subsequent steps of the analysis to capture shape variation independent of allometric effects of size. The vector of shape change between high and low altitude parental populations was calculated from the difference between the mean shape for the three highland and three lowland strains used as parents in F20 crosses. Shape data (represented by the joint contribution of PC2-26 or PC2-58) from individual wings were projected onto this difference vector to create a ‘shape score’ that aligns with the primary axes of non-allometric shape variation between the divergent populations.

### Genomic Sequencing of samples

For a given F20 cross and sex, we phenotyped at least 1100 individuals (see Supplemental file 2 for sample sizes), and combinatorically selected the 50 most extreme individuals from both the distribution of shape (measured as a shape score, described below) and size (measured as centroid size). From each cross and sex, those individuals with the 50 highest and 50 lowest shape scores were selected for sequencing while wing size pools were created from the individuals with the 50 largest and 50 smallest wings. In cases where an individual was an outlier for both size and shape, these individuals were included in both pools (details below, Figures 2, S4).

**Figure 2.**
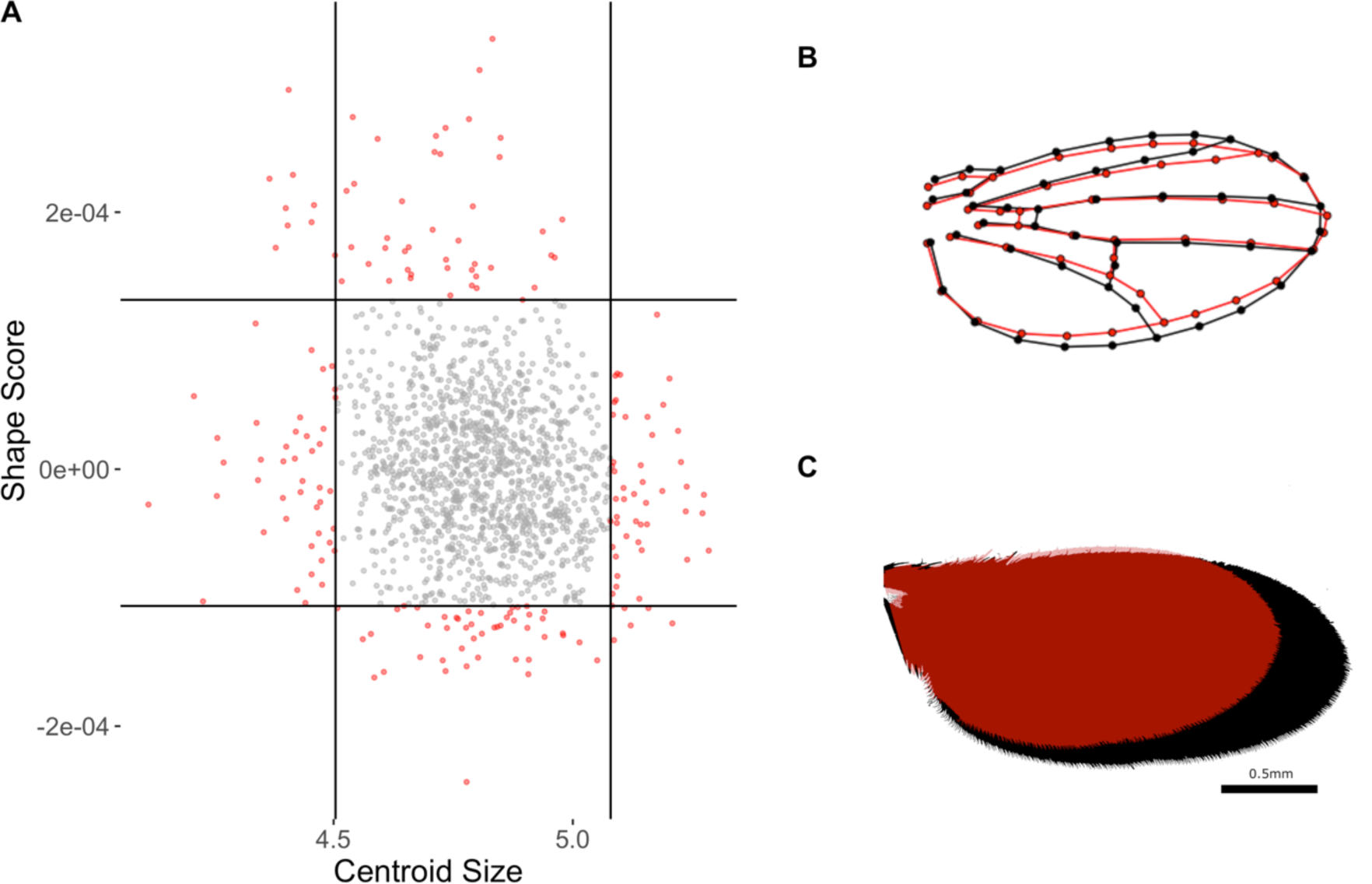
Selection of bulk pools used for sequencing from one F20 population. (A) Distribution of individual (each dot) phenotypes along the size and shape score axes. The 50 most extreme individuals along each axis were selected for sequencing and are indicated in red. Only males are shown because of the substantial size dimorphism. (B) Mean shape difference between individuals in the extreme pools along the shape score axis. Shape change is magnified 2x for visualization. (C) Mean size difference between pools selected for sequencing.

Genomic DNA extractions were performed using a Qiagen DNeasy kit on a maximum of 25 individuals in each sample. To create pools for sequencing, samples were combined such that each extraction sample of flies contributed an equimolar amount of DNA to the sequenced pool, using concentrations measured with the DeNovix dsDNA Broad Range Kit. The number of flies in each extraction sample and the amount of DNA added to the final pool can be found in supplemental file 1. Library prep using the NEB Ultra II kit for library preparation and 150bp paired end sequencing using illumina NovaSeq6000 was performed by Genome Quebec (Montreal, QC, Canada).

### Genomic analysis

Reads were trimmed with Trimmomatic (v0.36) to remove adapter contamination and regions of low quality, and checked for using FastQC prior to alignment (Bolger et al., 2014). Trimmed reads were aligned to the *Drosophila melanogaster* genome (v6.23) using BWA-MEM (v0.7.8) (Li and Durbin, 2010). Sequencing replicates of the same biological samples were merged using SAMtools (v1.11). PCR duplicates were removed using Picard with the MarkDuplicates tool (v 2.10.3) and reads with a mapping quality score less than 20 were removed using SAMtools (Li et al., 2009). A local realignment around indels was performed using GATK using the IndelRealigner tool (v3.4.46). Highly repetitive regions of the *Drosophila* genome were identified and subsequently masked in mpileup files using RepeatMasker (v4.1.1) with default settings. INDELs and regions within 5bp of an indel were identified, and masked, using PoPoolation2 scripts (v1.201) (Kofler et al., 2011b). F_ST_ was calculated in 5,000bp windows using grendalf (v0.2.0) (Czech et al., 2023). F_ST_ values that were calculated to be negative after correcting for sampling variance, were considered as a value of 0 for plotting and analysis. To identify differentiated loci shared between all three sequenced crosses, a modified Cochran-Mantel-Haenszel (CMH) test was used. Sampling effects were accounted for using the ACER package (v.1.0) in R, using the number of phenotyped individuals for each cross as population size, with 0 generations of divergence (drift) between selected pools (Spitzer et al., 2020). To adjust for multiple testing, we used the Benjamini-Hochberg correction (Benjamini and Hochberg, 1995) with an adjusted alpha of 0.05. BEDtools (v2.19.1) was used to identifiy the nearest gene to each site passing the significance filter (Quinlan and Hall, 2010). We performed a GO analysis for the set of genes with significant SNPs annotated within 2000 bp using Gowinda, with 100000 simulations (Kofler and Schlötterer, 2012).

### Functional testing using RNAi of candidate

To estimate the effects of candidate gene knockdown on shape change, we focused on genes in regions of chromosome 3R for the shape QTL (3R:22,580,970..22,733,819). We expressed an RNAi construct for the **G**ene **O**f **I**nterest (GOI) in the developing wing imaginal disc. RNAi experiments were done using fly media with the Dworkin lab recipe (see supplementary file 1). A nubbin-Gal4 line (BDSC:25754; genotype: *P{w[+mC]=UAS-Dcr-2.D}1*, *w[1118]*; *P{w[+mW.hs]=GawB}nubbin-AC-62*), carrying a driver known to express broadly in the developing wing pouch and widely used for the study of wing development (example: (López-Varea et al., 2021), was crossed to RNAi lines (UAS-GOI.RNAi) and progeny were allowed to develop at 24°C at low larval density. All but one RNAi line were derived from the *Drosophila* transgenic RNAi project (TRiP) panel (Perkins et al., 2015) and a common background with the TRiP control (BDSC:36303; genotype: *y[1] v[1]*; *P{y[+t7.7]=CaryP}*attP2) was used in control crosses (to the *nubbin*-Gal4). The genes for which RNAi lines were used in the experiment were: *locomotion defects* (*loco*), *winged-eye* (*wge*), *button* (*btn*), *wake*, *eukaryotic translation elongation factor alpha 2a* (*efa6*) and *TGFß-activated kinase 1-like2* (*takl2*) (Table S1). The *loco* RNAi line also carries a spontaneous *scute* allele. Given that *scute* influences aspects of bristle development on the wing, it may have effects on wing shape that are currently unknown. To account for this effect, *loco* RNAi flies were also crossed to a *white (w[1118])* mutant bearing strain, co-isogenic to the GAL4 line, to assess the effects of the strain independent of *loco* knockdown.. The genetic background for *wge* RNAi line was unknown, so we crossed this line to a *w[1118]* mutant strain as well, as an additional control for this construct. Each cross was done in two replicate vials. Adult F1 flies were collected 24-48 hours following eclosion, to allow for wing hardening, and stored in 70% EtOH until dissected and imaged for morphometrics. Shape data were collected from ∼35 individuals of each sex for each cross using the 15-landmark method (Supplemental File 2).

The effect of RNAi knockdown on shape was estimated by fitting a model with the procD.lm function (geomorph package) with predictors for log centroid size, genotype of the cross and the interaction of the two as well as a term for the replicate vial as a source of error. Males and females were considered separately in this analysis, as the altitudinal effect vectors were also calculated separately. Least squared means for each cross genotype were estimated, and contrasts between RNAi crosses and the appropriate control (Table RNAiLines) were calculated to generate the effect vector for RNAi knock down for plotting shape changes. Pairs bootstrapping was used to estimate distributions of vector correlations between altitudinal difference vector and the relevant RNAi knockdown effect vectors. Then the RNAi effect vector was calculated for each RNAi knockdown and the appropriate control from the same model as was used in the geomorph analysis (the effects of centroid size, genotype and their interaction plus the effect of replicate vial within genotype). Observations were sampled with replacement, in a non-stratified manner (sample sizes for groups were approximately equal), for 1,000 iterations to generate distributions of bootstrapped values.

### Quantitative complementation mapping of candidate genes on chromosome 3R

A genomic deletion series from both the *Drosophila* Deletion (DrosDel) (Ryder et al., 2007) and Exelixis (Parks et al., 2004) genomic deletion panels, spanning the region of differentiation on chromosome 3R (22,580,970..22,733,819) in the ZI192N x EF96N and ZI192N x EF81N crosses were used for quantitative complementation mapping of the contributing allele. A full list of the deletion lines used, and the genes predicted to be included in deleted regions, is included in supplementary file 2. In addition to the lines used in mapping crosses, we included a number of other high and low altitude lines: EF 43N, 81N, 96N and 119N ; ZI 192N, 251N, 360N, 357N, to test for effects at the population level. Virgin females from each of the Ethiopian and Zambian derived strains, were crossed to males bearing the genomic deletion, and allowed to lay eggs for 2 days on Dworkin lab fly medium (Supplemental file 1). Each cross was done in two replicate vials. F1 heterozygotes (for the deletion) were selected based on genetic markers demonstrating absence of the balancer chromosome, and preserved in 70% ethanol until males were dissected and imaged following the protocol outlined above. Shape data were collected using the 15-landmark method. Each deletion panel was considered separately (as they were generated in different isogenic wild-type strains, and had distinct control strains) for statistical analysis but using the same methods. A mixed model, accounting for the influence of size, deletion line, and source population, all two-way interactions between these terms plus a random effect for the replicate vial nested within cross and for genetic lines within background (shapescore ∼ (CS+delLine+background)^2 + (1 | delLine:background:rep)) was used to test associations with shape change. The response variable was the shape score calculated by projecting superimposed shape data onto the vector based on the mean difference between high and low altitude populations.

### Patterns of allometric variation in wing shape across the F20 crosses

For all morphometric analyses, male wing shape data from three high altitude (EF43N, EF81N, EF96N) strains, three low altitude (ZI192N, ZI251N, ZI418N) strains, and 3 crosses (ZI192N x EF43N, ZI192N x EF81N, ZI192N x EF96N) were used. Each distinct group, whether an inbred strain or F20 intercross is referred to as a ‘genotype’ throughout the analysis for convenience. All data for these analyses were collected using the 15-landmark method and jointly superimposed.

We examined variation in allometric (shape-size) relationship across genotypes, within and between populations. Allometric effect vectors were estimated using a model regressing wing size, genotype, and their interactions on shape. To aid in visualization of allometric effects, we projected observed landmark configurations onto the estimated allometry vectors (shape score). Similarly, we projected predicted values along these vectors. These were calculated with the plotAllometry function from the geomorph package. Using the pairwise() function within RRPP/geomoph, vector correlations among allometric vectors were calculated at the strain level. Additionally, to understand group level changes (high altitude lines, low altitude lines and F20 crosses), the effect vectors within group were estimated from a linear model regressing the effect of wing size, population, and the interaction of the two on shape. Confidence intervals for both statistics were estimated using non-parametric bootstrapping of observation vectors.

To compare the selection vector used in this experiment with the direction of shape variation in the population, mean shape from 10 additional Zambian and 11 additional Ethiopian strains was obtained from Pesevski and Dworkin (2021). One strain was shared between studies to allow for the estimation of effects between the two data sets. An approximate G matrix could then be calculated using a PCA of the line means of the 6 strains used in this study plus the additional lines. Shape change vectors were calculated by finding the difference between mean shape vectors for the Zambian and Ethiopian populations. The effect of lab rearing conditions, landmark acquisition by different researchers along with wing centroid size were accounted in the linear model.

## Results

### Size and shape adaptation have a polygenic basis

QTLs contributing to wing size variation were mapped by measuring genetic differentiation between large and small pools using F_ST_ in 5,000bp windowed analysis. Overall, the genetic basis of size adaptation to high altitude appears to be polygenic, with no striking regions that would indicate alleles of large effect contributing (Figure 3). For further analysis, windows with an F_ST_ in the top 1% of all values were included. This selected cutoff is somewhat arbitrary but no less so than for a specific significance threshold. A small number of outlier windows are shared between different crosses (total = ZI192N x EF96N: 254, ZI192N x EF81N: 253, ZI418N x EF43N: 262). Two outlier windows were “shared” between ZI192N x EF91N and ZI192N x EF81N, 4 shared between ZI192N x EF96N and ZI418N x EF43N and 3 shared between ZI192N x EF81N and ZI418N x EF43N. Shared windows could indicate either a shared causative allele contributing to shape variation in both crosses or could reflect chance linkage disequilibrium of genomic regions linked to different causative alleles. When all populations are considered together using a modified CMH test which accounts for sampling, 45,802 SNPs, representing a total of 2,769 unique genes, are identified as significant using a FDR threshold of 0.01 (Figure S5, Supplemental file 3). Most of these associations can be explained though linkage to a subset of causative SNPs, yet despite this, it is clear this represents a highly polygenic genetic architecture. GO analysis of the genes nearest to the annotated SNPs, did not identify any enriched terms.

**Figure 3.**
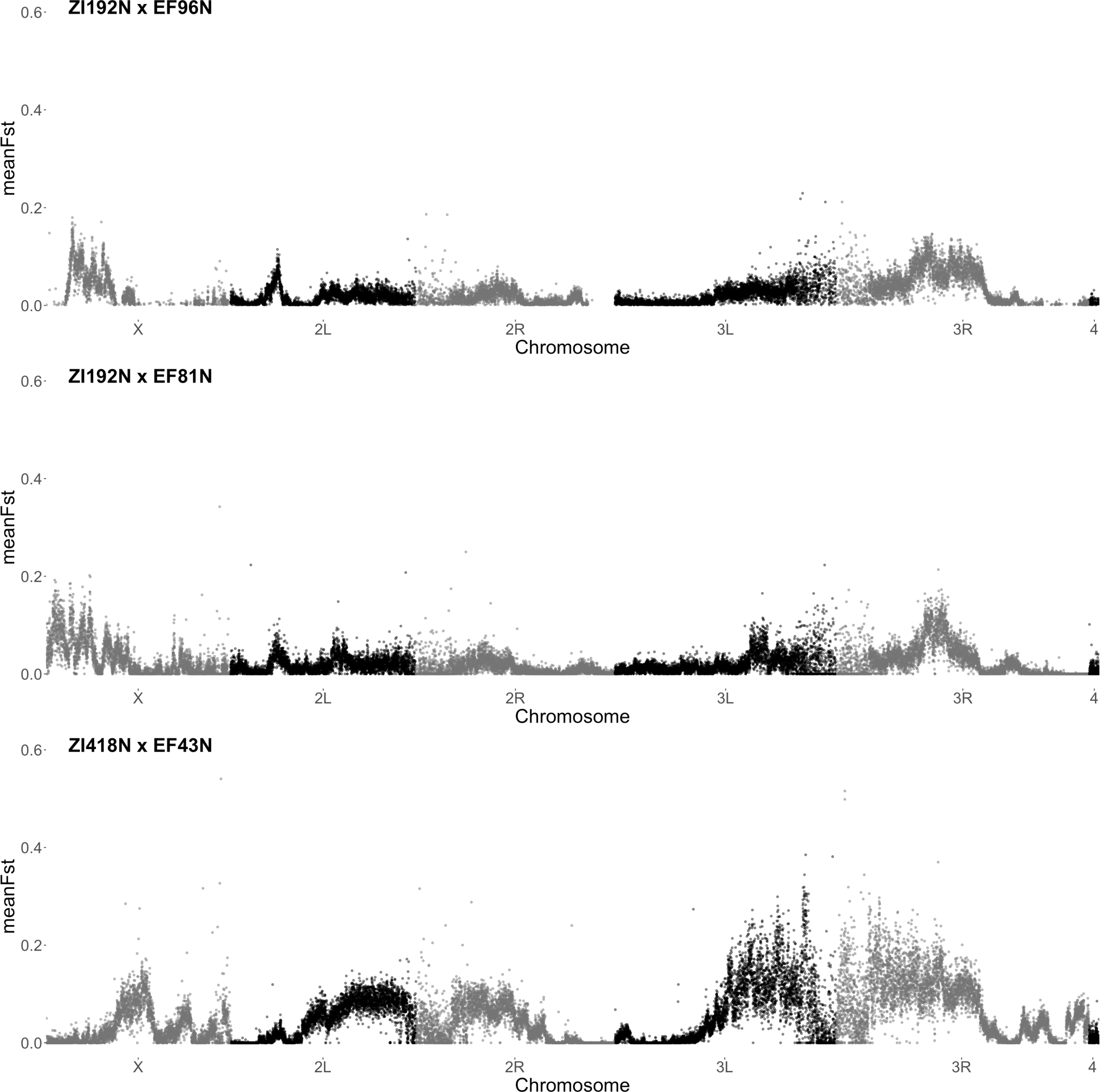
Genetic differentiation indicates a polygenic basis of size divergence. F_ST_ between pools of individuals for mapping (males), in 5000 bp windowed analysis using grendalf (v0.2.0)(Czech et al., 2023). Parental genotypes of crosses are indicated above each panel.

In contrast to size, mapping changes in shape between high and lowland populations revealed specific regions of the genome contributing to the shape variation (Figure 4). Although a large region of differentiation is shared on chromosome 3R in both the ZI192N x EF96N and ZI192N x EF81N crosses, this region is not differentiated in the ZI418N x EF43N cross (Figure 4, S6). There is a large region of differentiation on chromosome 2R in the ZI418N x EF43N cross that appears to be unique to this cross. No outlier windows, using the top 1% of F_ST_ values cutoff, are shared between ZI418N x EF43N and other crosses (total = ZI192N x EF96N: 254, ZI192N x EF81N: 253, ZI418N x EF43N: 262). There are 141 windows shared between ZI192N x EF96N and ZI192N x EF81N, with all but one of those windows on chromosome 3R between 3R:21,455,001 – 23,275,001 (Figure S6). The large genomic regions contributing to shape differentiation make it difficult to identify specific alleles that may be contributing in these regions. When all crosses are considered together using the modified CMH test, the large region on chromosome 3R is identified as substantial (Figure 5) due to the effect in two of the three crosses (Figure 4). Using this test, a total of 20,224 SNPs, in 6,395 genes, are identified as having significant associations with shape adaptation (Supplemental file 5). Again, most of these sites are due to LD with causative sites. A GO analysis of genes found within QTLs found no significantly associated terms.

**Figure 4.**
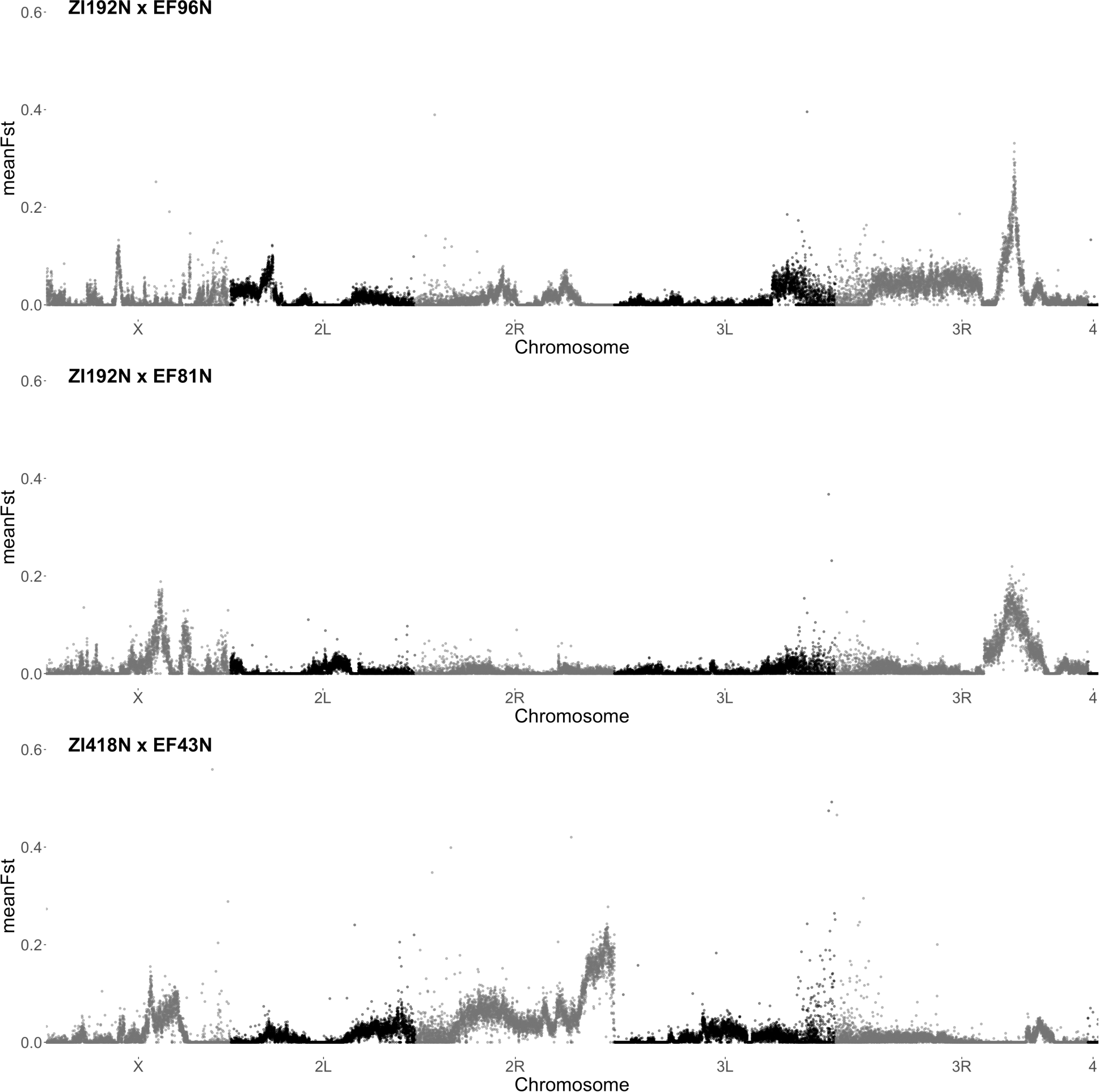
Genetic differentiation indicates a polygenic basis of shape divergence. F_ST_ between male pools measured in a 5000 bp windowed analysis using grendalf (v0.2.0)(Czech et al., 2023). Parental genotypes of crosses are indicated above each panel. Note that for the crosses in the first two rows, with a shared Zambian parental strain, there is a shared peak of divergence on chromosome 3R.

**Figure 5.**
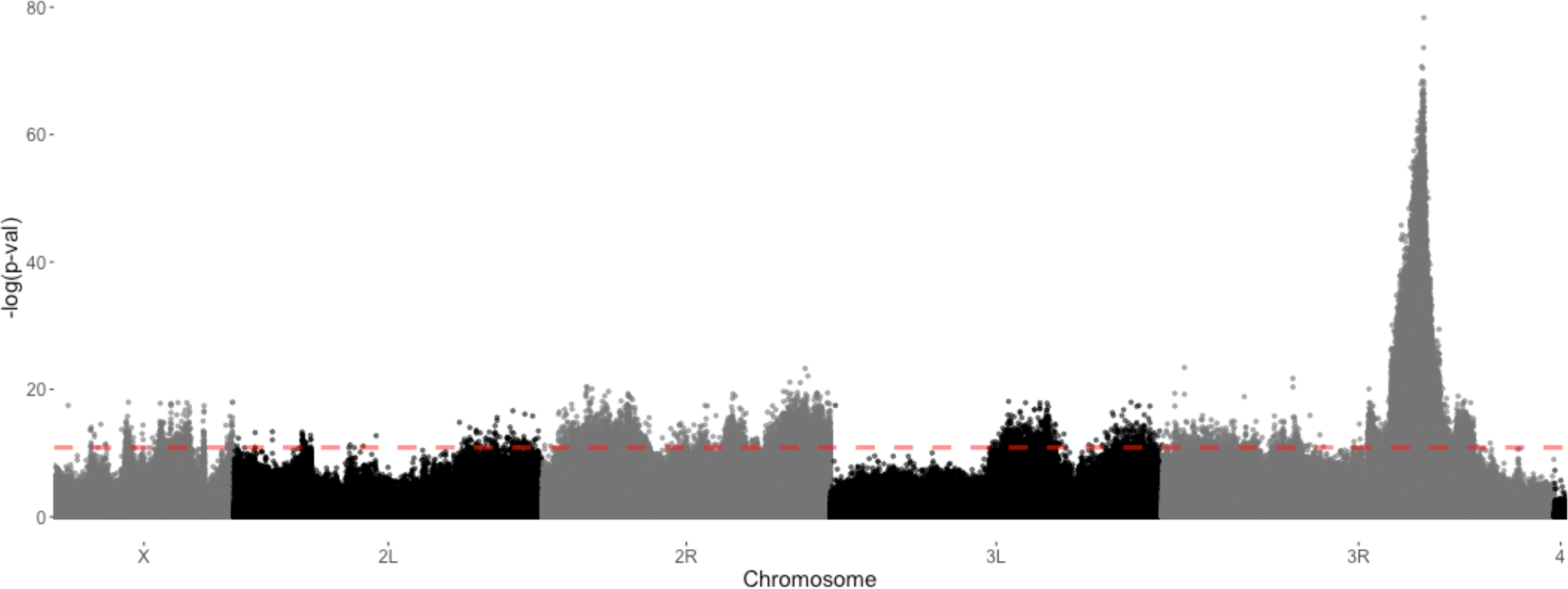
CMH test for association of SNPs with shape divergence between high- and low-land populations in males, using data from all crosses. Modified CMH test to account for pooled sequencing sampling error, as implemented in ACER (Spitzer *et al* 2020), using population sizes equal to the total number of individuals phenotyped for each cross and 0 generations of selection between pools. Red line indicates adjusted FDR of 0.01.

There is also evidence for sex specific alleles contributing to shape differentiation (Figure S7). Sexual shape and size dimorphism have been documented for the *Drosophila* wing in these populations (Pesevski and Dworkin, 2020). In the case of the QTL on 3R in the ZI192N x EF96N and ZI192N x EF81N crosses, this region may contribute more in females as the allelic differentiation in this region is greater. In the ZI192N x EF81N cross regions of the X chromosome are differentiated only in the males,. This same pattern on the X chromosome, with the addition of another region on chromosome 2R appearing to contribute more in one sex than the other. While these observations can be explained by sampling variance, it is more likely that there is varying expression of allelic effects between sexes.

### Evaluating candidate genes in the chromosome 3R genomic region contributing to shape divergence

Candidate genes located in the QTL region on chromosome 3R that might influence wing shape were selected for follow-up analysis. While RNAi-mediated knockdown of one gene, *loco*, had a substantial effect on wing shape when compared to controls (Figure S8), given that the mutational target size of wing shape is large, it is expected that perturbation of many genes are likely to show effects. As such, simply identifying significant genes influencing wing shape is unlikely to be informative with respect to which genes may be contributing to the divergence in wing shape between the low and high-altitude populations. A far more informative and relevant approach is to investigate the degree of a shared directions of effects of gene knockdown with the altitudinal effect vector, in terms of wing shape. For all 6 examined genes, the correlation between RNAi knockdown vectors with the altitudinal divergence vector was moderate (Figure 6). The *wge* RNAi vector shows the largest observed correlation (0.57; CI: 0.34 – 0.65), suggesting a moderately shared direction of effect with altitudinal divergence. However, other genes showed similar levels of correlation as well: *loco* (0.46; CI: 0.20 – 0.62), *ef6a* (0.53, CI: 0.35 – 0.67), and *takl2* (−0.39; CI: −0.57 - −0.16) with the altitudinal effect vector. Shape changes these three genes are like that of the altitudinal effect vector, and thus aligned with the direction of adaption, making them good candidates for future work (Figures 6, S9). Interestingly, in most cases, there is relatively low correlation (low shared direction of effects) between RNAi knockdown shape change vectors themselves (Table S1).

**Figure 6.**
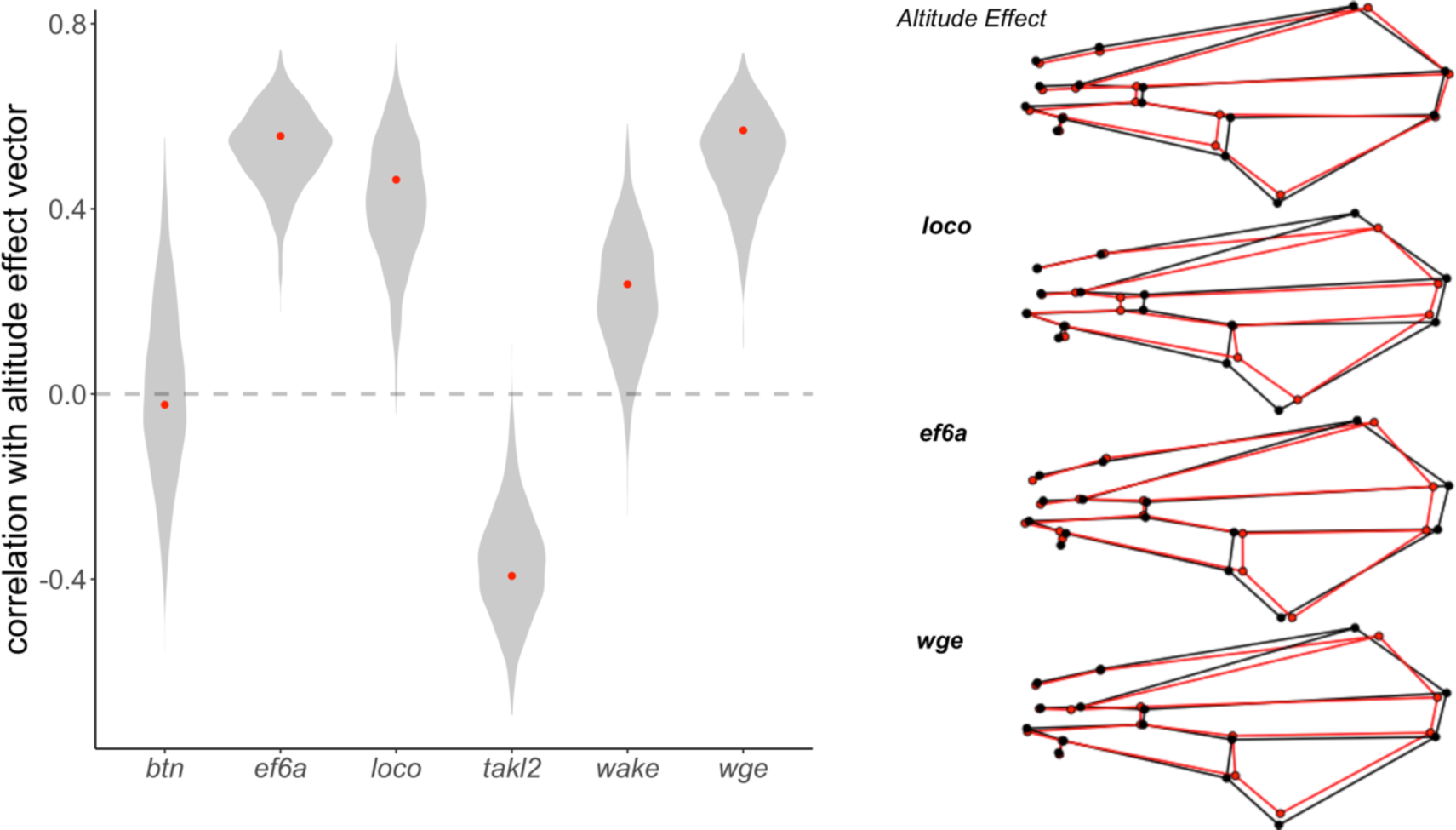
RNA interference knockdown effects of genes in the chromosome 3R candidate region. Bootstrapped distribution of correlations between altitudinal effect vectors, and RNA interference knockdown vectors, with observed value noted by red points. Wing shape changes plotted for altitudinal effect and selected RNAi effect vectors. For altitudinal effect vector, black represents the low altitude and red the high altitude shape. For RNAi effect vectors, black represents controls and red, the RNAi knockdown. Shape change effects have been magnified for visualization: altitudinal effect 20x; *loco* effect: 1.5x, *ef6a* effect: 2x, *wge* effect: 3x.

### Quantitative complementation analysis of candidate genomic region

In addition to candidate gene knockdowns, we also used co-isogenic deletion mapping strains to perform quantitative non-complementation tests between smaller genomic regions and shape change to identify smaller regions within the QTL that contribute to the shape change of interest. As shown with the interactions plots, there was little evidence for substantial effects of population of origin for most tested deletions (Figure 7). For both panels tested, there was no substantial effect of the interaction between population and deletion line (Tables S3, S4). In the DrosDel panel, there may be different reaction norms between the two populations for the 20515 line (Figure7). This line is predicted to have no deletions in genes that overlap with the previously identified candidates (See supplementary file 7 for full list). In the Exelexis panel, deletion line 7671 is a possible candidate genomic region containing variants influencing shape (Figure 7). This region contains both *wake* and *wge*. Additionally, deletion 7740 in the Exelexis panel may be a candidate, and contains the genes *loco*, *btn* and *efa6* from the candidate list.

**Figure 7.**
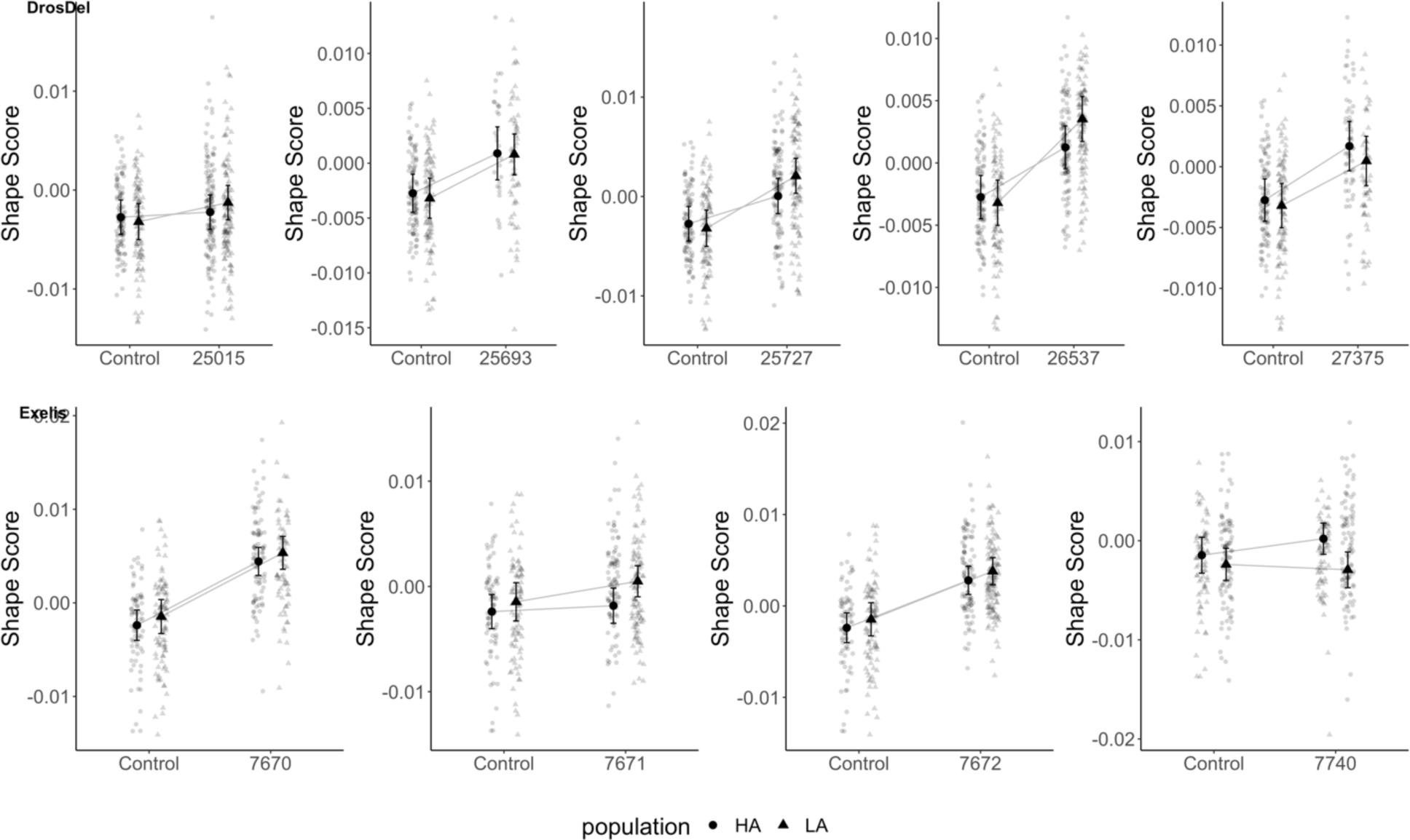
Quantitative complementation mapping. Estimated shape scores, calculated from projection onto altitudinal effect vector, between control and deletion cross. 4 high altitude and 4 low altitude lines were used to estimate population means for high altitude (triangles) and low altitude (circles) populations. Grey points indicate observed data and black points indicate estimated means. Error bars represent 95% confidence intervals.

### Size and shape divergence have partially independent genetic bases

Because size and shape are tightly linked though allometry, we wanted to ask if there was a change to allometric effect vectors in addition to the differences in shape between crosses in this study. As environmental variation was tightly controlled in these experiments, the majority of phenotypic variation we observe should be genetic. Wing size is intermediate in the F20 cross genotypes compared to the parents (Figure S10). Using three crosses with a shared Zambian parent but unique Ethiopian parents, we estimated allometric effect vectors to compare the relationship between size and shape in crosses compared to parental genotypes. There is a significant effect of allometry between genotypes; however, this only explains a small proportion of overall shape variance (*R^2^ = 0.0013, F = 4.36, p < 0.001*, Table S5). Despite different allometric effects within Ethiopian parental strains and between Ethiopian and Zambian parents, we observe a shared direction of allometry in all three F20 crosses (Figure 8A, Table 1). However, the overall correlation of allometric effect vectors between the cross and parental populations remains about the same (Figure S11). It is important to note the larger magnitude of allometric effect vectors in parental populations (ZI: 0.22, CI: 0.15 – 0.27; EF: 0.11, CI: 0.11 – 0.20) compared to the cross populations (0.06, CI: 0.06 – 0.07)(Figure S11). Most of this variation is related to a shift in the position of the L2 and L4 longitudinal veins and a posterior shift in the anterior cross vein between the smallest and largest files (Figure 8C). There is a high degree of individual variation within both the crosses and the parental lines (Figure 8B).

**Figure 8.**
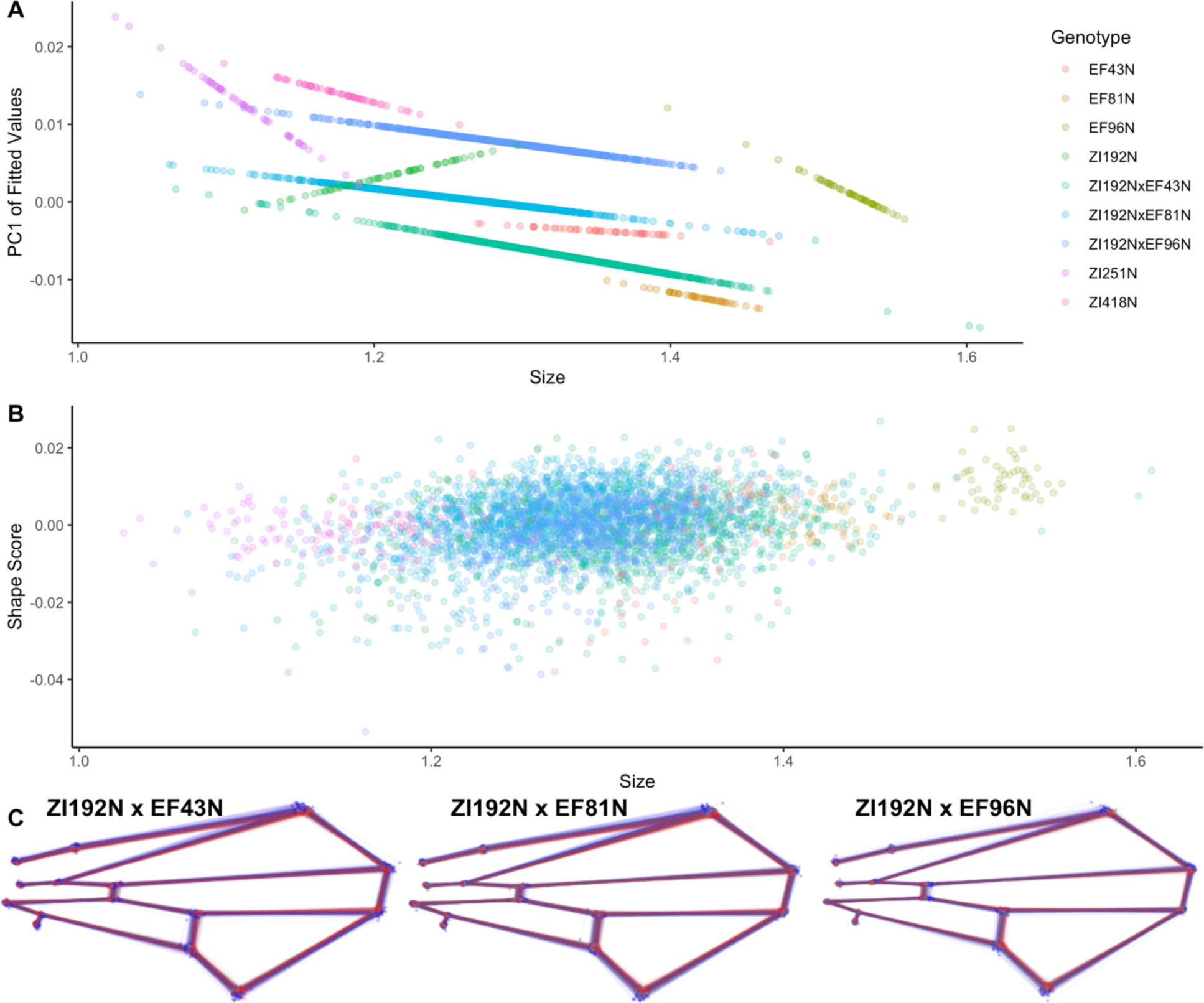
Similar allometric component of shape in F20 intercross despite unique relationship in parental lines. (A) Model predicted allometry is represented by slope of lines. PC1 of model fitted shape residuals plotted against wing size. (B) Shape scores of individuals calculated by projection of individual shape configurations onto the allometric effect vectors for each genotype to represent within line variation of allometric relationship. (C) Wire frames aid in visualization of shared allometric relationship across the three distinct F20 crosses. Diversity of shape variation is plotted in grey, with the largest (blue) and smallest (red) 10% of wings highlighted for each cross.

**Table 1.** Pairwise vector correlations between estimated slopes of allometric effects. 95% CI (via bootstrap) for estimates are indicated in brackets.

|  | EF43 | EF81 | EF96 | ZI192 | Zi192Ef43 | Zi192Ef81 | Zi192Ef96 |
| --- | --- | --- | --- | --- | --- | --- | --- |
| EF43 |  | 0.23<br>(-0.42, 0.58) | 0.25<br>(-0.06, 0.44) | 0.90<br>(0.77, 0.90) | 0.71<br>(-0.14, 0.95) | 0.90<br>(0.05, 0.98) | 0.79<br>(0.05, 0.96) |
| EF81 |  |  | 0.54<br>(-0.15, 0.62) | 0.13<br>(-0.41, 0.51) | 0.40<br>(-0.31, 0.57) | 0.39<br>(-0.33, 0.58) | 0.37<br>(-0.32, 0.56) |
| EF96 |  |  |  | 0.13<br>(-0.17, 0.35) | 0.67<br>(0.06, 0.73) | 0.58<br>(-0.02, 0.72) | 0.58<br>(0.02, 0.70) |
| ZI192 |  |  |  |  | 0.62<br>(-0.42, 0.87) | 0.82<br>(-0.27, 0.90) | 0.77<br>(-0.19, 0.91) |
| Zi192Ef43 |  |  |  |  |  | 0.88<br>(0.83, 0.99) | 0.82<br>(0.72, 0.97) |
| Zi192Ef81 |  |  |  |  |  |  | 0.91<br>(0.83, 0.98) |
| Zi192Ef96 |  |  |  |  |  |  |  |

The selection vector used to score wings in the F20 mapping crosses was defined as the difference vector between the three Ethiopian and Zambian lines used for the mapping populations. Because these lines were chosen at random regarding size and shape diversity, this vector does not necessarily provide a perfect representation of shape differences between highland and lowland populations. To address this, we calculated the same shape change vector using additional data from 11 Ethiopian and 10 Zambian lines. The shared direction between difference (altitudinal effect) vectors to the direction of genetic variation in the population, using the approximate **G** matrix, was measured by projecting the mean shape of each line onto the difference vector to calculate a shape score and then calculating the relationship between these vectors and the axis of variation, summarized in PCs. There is a stronger relationship between the direction of greatest variation among strains (PC1) and the altitudinal effect vector using the subset of lines selected for this study (r = 0.61) than the total population (r = 0.22)(Figure S12). There is also a relationship between both effect vectors and PC2 (subset: r = 0.54; all data: r = 0.52).

## Discussion

The main goal of this study was to use experimental mapping populations to estimate aspects of the genetic architecture influencing wing shape and size contributing to adaptation to a high-altitude environment. We demonstrate a polygenic architecture of adaption for both traits, based on mapping of three F20 crosses (Figure 3,4). Despite the intrinsic link between shape and size of traits though shape-size allometry, we demonstrated that the alleles contributing to shape divergence and size divergence could be identified independently (Figure 3,4). Additionally, despite different allometric shape-size relationships in parental inbred lines, F20 populations from the crosses had similar shape-size allometries, indicating that the alleles contributing to shape and size variation independently may not affect the allometric relationship between the two (Figure 8). Overall, this work provides support for a model in which modest (or at most moderate) shifts in allele frequency for many genes, rather than fixed differences among alleles for a small number of genes, contributes to adaptation for the traits of interest.

Wild populations of *D. melanogaster* have large effective population sizes, with high genetic diversity (Duchen et al., 2013; Sprengelmeyer et al., 2020). As wing size and shape both have large mutational target sizes (Carreira et al., 2013, 2011; Houle and Fierst, 2013; Weber et al., 2005, 2001), it is likely that the ancestral population was already segregating considerable “relevant” genetic variation that selection could act upon, ultimately contributing to adaptative divergence. As such the relative contribution of new (*de novo*) mutations concurrent with selection will likely play a more modest role in adaptation in these populations. In the *Drosophila* Genetic Reference Panel (DGRP), SNPs with replicable effects on wing size were identified in over 30 genes (Vonesch et al., 2016). For shape, over 500 SNPs contributing to variation were identified in the DGRP (Pitchers et al., 2019). In both cases, this is likely an underestimate of the number of genetic variants contributing to trait variation, as polymorphisms with very small effect sizes, in aggregate can have substantial effects on the variation of complex traits, yet remain difficult to identify with GWAS methods (Boyle et al., 2017; Rockman, 2012). Given the large effective population size and large mutational target of the traits studied, it is perhaps not surprising that this work identified a polygenic basis of adaptation without evidence of alleles of large genetic effects that were fixed between populations. This same polygenic pattern of rapid adaptation following an optimum shift has been previously reported using both simulations (Jain and Stephan, 2017), mapping of life history traits in guppies (Whiting et al., 2022) and craniofacial shape between mouse species (Pallares et al., 2016). Small phenotypic effects of alleles and large effective population size, as seen in the case of the *Drosophila* wing, are two predictors of rapid adaptation with many alleles contributing (MacPherson and Nuismer, 2017).

In contrast to the polygenic architecture of wing size variation identified in this study, previous work has identified some genes harbouring alleles of moderate effect sizes contributing to size variation along latitudinal clines. Small intronic deletions were identified as associated with clinal variation on in both North American and Australian populations as allele frequencies varied along the cline (Paaby et al., 2010). GWAS studies have identified a number of other alleles within the insulin signaling network as associated with clinal size variation (Fabian et al., 2012) with a particularly strong candidate being a SNP in *foxo*, accounting for as much ∼16% of wing size variation in the samples from the North American populations (Durmaz et al., 2019). In contrast, both this study (Figure 3) and a previous similar study (Sprengelmeyer et al., 2022) using these altitudinally diverged populations identify a polygenic architecture of adaptation but not an enrichment for genes associated with insulin signaling. In both the case of altitudinal and latitudinal variation, *Drosophila* populations have large effective population sizes with a high degree of segregating variation. Why is it that a single genetic pathway seems to contribute more in one case than the other? It is possible that other factors such as the significant contribution of genomic rearrangements (Kapun et al., 2016; Kennington et al., 2007; Rako et al., 2006; Weeks et al., 2002) and highly pleiotropic nature of variants (Durmaz et al., 2019; Paaby et al., 2014), contributing to latitudinal variation could in part explain these differences.

One QTL with a significant contribution to shape adaptation was identified on chromosome 3R (Figure 4,5), but was only identified in two crosses with a shared Zambian parental strain. While there is a large effect allele in this region, it is not fixed in the population as the ZI418N x EF43N cross did not have allelic differentiation in this region as would be expected if the allele was fixed. This highlights the importance of using a number of distinct genetic backgrounds for crosses when considering the alleles contributing to adaptive divergence in wild populations. Many of the classic examples of adaptation with the majority of variance explained by single large effect alleles mapped these alleles using a limited number of genetic backgrounds. For example, the identification of large effect alleles explaining differences in coat colour in beach mice (Steiner et al., 2007) or variation in pelvic spine size in sticklebacks (Shapiro et al., 2004) required only the genotyping of hundreds of F2 individuals from a single genetic cross. In some cases, including more backgrounds and crosses has supported this finding of large effect alleles being fixed or near fixed in populations and explaining the majority of divergence, as is the case of alleles for *ectodysplasin* contributing to stickleback armored plate variation (Colosimo et al., 2005). However, more sequencing, larger sample sizes and more backgrounds used in mapping has revealed many additional loci that also contribute in this case (Miller et al., 2014). Although many of the large effect variants identified are important for adaptation, the sampling designs used in these studies is biased towards identifying larger effect alleles that can be detected with modest sample sizes. By increasing the number of independent crosses in this study, we can identify a more complex architecture of adaptation for both traits.

Although not fixed in the population, the QTL on chromosome 3R contains alleles contributing to wing shape divergence. In follow up experiments, we attempted to identify the gene(s) contributing to shape variation between high and low altitude populations in this genomic region. Using both RNAi of candidate genes and quantitative complementation mapping using genomic deletions in this region (Figure 6,7), we did not identify a single ‘standout’ candidate, instead identifying multiple genes with shape effects, each moderately aligned with the direction of altitudinal divergence. The most likely explanation is that multiple genes harbour alleles jointly contributing to phenotypic differences. Clusters of alleles of small phenotypic effect, grouped together in LD and contributing to divergence, are predicted in the presence of gene flow (Yeaman, 2013) and demonstrated in empirical studies (Enbody et al., 2023; Orteu and Jiggins, 2020). Given the distinctly multidimensional nature of shape, our findings may suggest a slightly altered version of this phenomenon. We observed that RNAi-mediated knockdown of multiple genes in the shape QTL on chromosome 3R showed moderate alignment with the vector of shape divergence (Figure 6, SFig 9). However, the correlations of effects between the RNAi vectors themselves was generally low (Table S1).

Assuming that effects of the segregating alleles in the genes is similar to that of their RNAi knockdown vectors, then it may be the aggregate direction of effects that we have mapped, and thus is possibly contributing to adaptive divergence in shape. Alternatively, it is possible that these effect vectors do not estimate directions of effects well, and further titrating the strength of knockdowns may help to obtain better estimates. It is also possible that the RNAi knockdown does not correlate with the segregating allelic effect of the candidate genes, although previous studies have demonstrated correlation between segregating polymorphisms and genetic knockdown effects for wing shape (Pelletier et al., 2023; Pitchers et al., 2019). Quantitative complementation mapping experiments identified two regions that may be related to shape change in the same direction as the altitudinal effect vector (Figure 7). One of these regions overlaps with *wge*; the same gene that demonstrated the highest correlation with phenotypic divergence via RNAi (Figure 6). *wge* has been shown to be important for expression of *vestigial*, a key wing patterning gene, in developing wing tissue (Katsuyama et al., 2005). Although there is a candidate gene in this region, it is not likely “the” gene contributing to divergence between populations as our evidence is not consistent with one single large effect allele accounting for divergence. From this data, it is not possible to determine at what frequency the causative allele or alleles are segregating in the wild populations as these alleles could not be identified. Although non-allometric components of shape varies along an altitudinal cline (Pitchers et al., 2013), and we accounted for allometric shape variation in our mapping study, changes in size and shape may still be intrinsically linked because of a shared genetic basis of the two traits. For example, the Hippo developmental network is best known for its role in regulation of organ size, including the size of *Drosophila* wings. Variants in genes in the hippo signaling pathway influence wing shape variation, with correlated phenotypic effects (Pelletier et al., 2023; Pitchers et al., 2019). Interestingly, allele frequency changes across multiple hippo signaling loci following artificial selection for wing shape changes did not result in substantial wing size change (Pelletier et al., 2023). In this study, we also see evidence for independent genetic bases of wing size and wing shape change (Figures 3,4). We observe a consistent allometric size-shape relationship in F20 crosses (Figure 8), indicating that despite shape changes, the allometric. These results should be interrupted with caution as wing morphology is highly plastic, with large environmental contributions to variance (Carreira et al., 2009; Debat et al., 2009). That is, the environmental effects contributing to shape and size variation may simply be more similar in the crosses than in the parental lines, creating a common phenotypic variance. However, because we still observe the partially independent genetic basis of both size and shape variation, this is strong evidence that these traits have independent optima and the changes observed are due to selection on the trait and are not explained simply though selection on the other trait.

This study adds to the growing body of evidence that adaptive divergence does not require alleles of large effect or sweeps of alleles and can instead be explained by small allele frequency shifts across many loci. Additionally, this work highlights the importance of using multiple genetic backgrounds when mapping the ‘alleles of evolution’ as the consideration of more genetic diversity in mapping cross designs can reveal more a more complex genetic architecture than what is identified using only a single genetic background. Further mapping of specific loci contributing to divergence in these populations will be an interesting element in understanding the origin of these alleles, that is if they are segregating in lowland populations or if they are in fact de novo mutations arising concurrently with adaptation that have not fixed in high altitude populations. Although alleles of large effect are important for explaining divergence in some situations, the patten of alleles contributing to adaptive divergence predicted by the geometric model is not universal in all cases. Future studies should address what ecological, population genetic and other factors are most important for determining the pattern of alleles that are ‘captured’ by divergence.

## Supporting information

SuppFile1

SuppFile2

SuppFile4

SuppFile6

SuppFile7

SuppFile5

SuppFile3

## Author contributions

Study Conceptualization and funding: ID, JEP

Study Design: ID, JEP, KP

Crosses: YH, KP, MB

Dissections: KP, MDS, IPD, MB

DNA extraction: KP

Phenotyping: KP

Analysis: KP, ID

Manuscript drafting: KP, ID

Manuscript editing: KP, ID, JEP

Manuscript revisions:?

## Data and script access

Upon acceptance all data and scripts will be made available both on github (https://github.com/DworkinLab) as well as a static copy (with DOI) on either Figshare or DRYAD. All raw sequence data will be deposited in NCBI SRA.

## Acknowledgement and Funding

This work was funded by NSERC Discovery and Discovery Accelerator grants to ID.

---

**Table S1.**
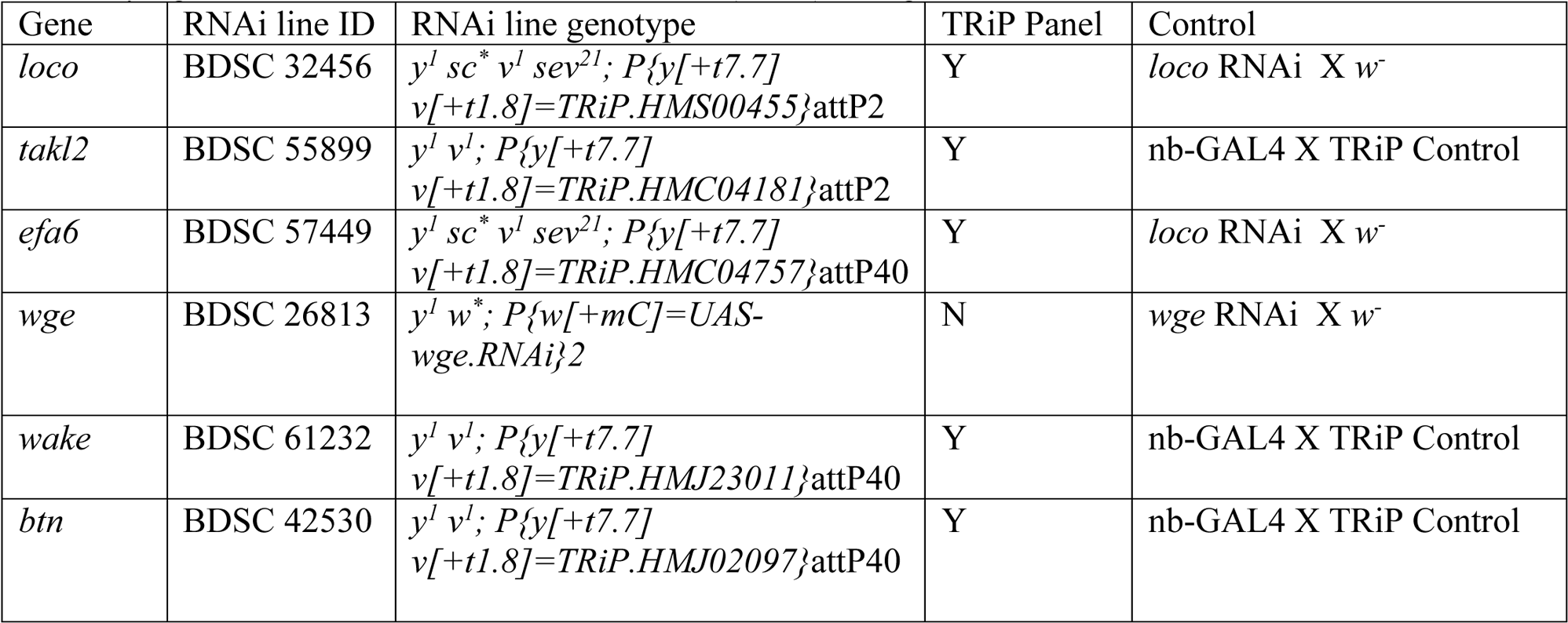
RNAi lines and appropriate controls for RNAi experiment. All experimental crosses were to the same *nubbin-Gal4* line (BDSC 25754) backrossed into the Samarkand background. In all cases, the TRiP control was (BDSC:36303), the third chromosome insertion line. *w^−^* were flies carrying the *white* mutation in the Samarkand (SAM) background.

**Table S2.** Pairwise correlations between estimated shape change vectors for RNAi knockdown. 95% confidence intervals for estimates are indicated in brackets.

|  | *loco* | *wge* | *btn* | *ef6a* | *takl2* | *wake* |
| --- | --- | --- | --- | --- | --- | --- |
| *loco* |  | 0.40<br>(0.19, 0.59) | 0.19<br>(-0.23, 0.54) | 0.050<br>(-0.19, 0.29) | -0.21<br>(-0.41, 0.04) | 0.23<br>(-0.03, 0.44) |
| *wge* |  |  | -0.27<br>(-0.61, 0.088) | -0.35<br>(-0.53, -0.16) | -0.25<br>(-0.45, -0.01) | 0.19<br>(-0.09, 0.40) |
| *btn* |  |  |  | 0.46<br>(0.19, 0.67) | 0.02<br>(-0.48, 0.39) | -0.10<br>(-0.59, 0.51) |
| *ef6a* |  |  |  |  | -0.27<br>(-0.41, -0.06) | -0.03<br>(-0.23, 0.20) |
| *takl2* |  |  |  |  |  | 0.23<br>(-0.03, 0.44) |
| *wake* |  |  |  |  |  |  |

**Table S3.** ANOVA table for the effect of deletion background from the DrosDel panel and wing size on wing shape.

| | $\chi^2$ | Df | Pr(>$\chi^2$) |
| --- | --- | --- | --- |
| Size | 50.02 | 1 | 1.53e-12 |
| Deletion Line | 115.22 | 5 | 2.2e-16 |
| Population | 0.88 | 1 | 0.35 |
| Size: Deletion line | 20.62 | 5 | 0.00095 |
| Size: Population | 7.81 | 1 | 0.0052 |
| Deletion line: Population | 9.80 | 5 | 0.081 |

**Table S4.**
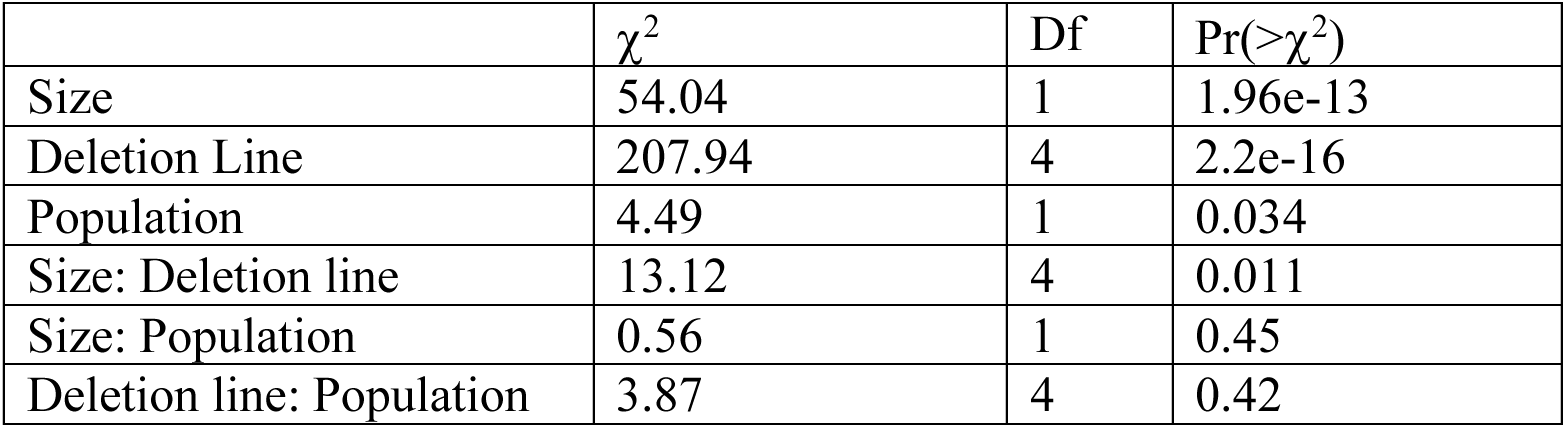
ANOVA table for the effect of deletion background from the Exelixis panel and wing size on wing shape.

**Table S5.**
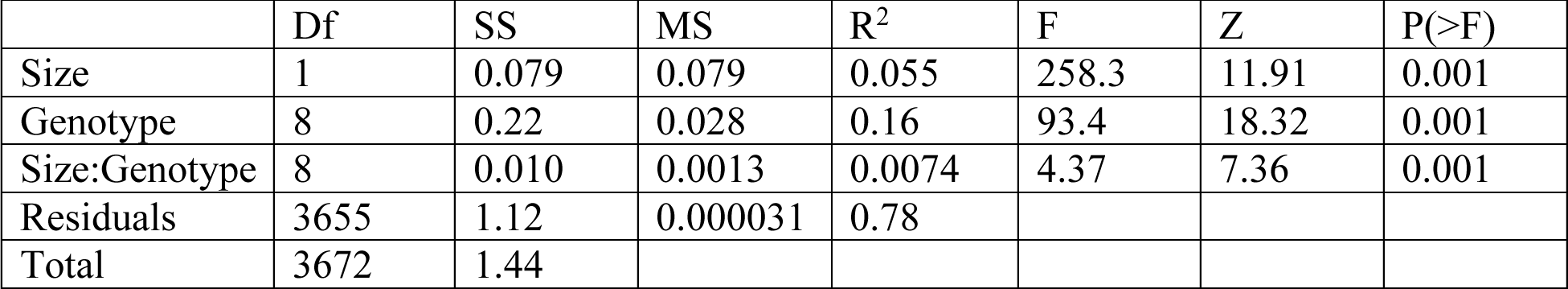
ANOVA table for the effect of genotype, size and the interaction on shape for the F20 cross genotypes compared to parental inbred lines. Model fit using residual resampling permutation test using RRPP/Geomorph.

**Figure S1.**
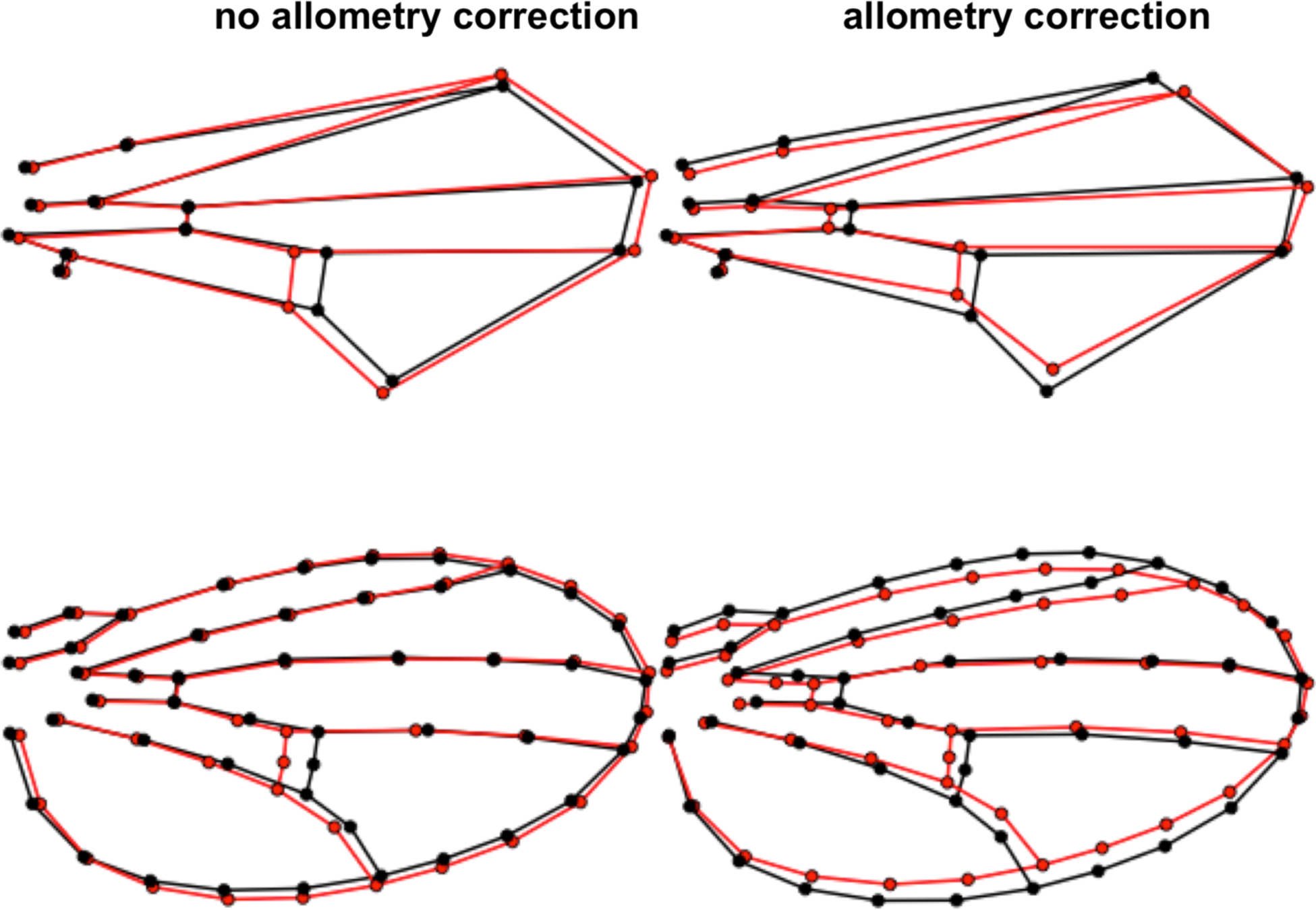
Shape change between high and low altitude populations is equivalent between methods. 15 point (top) and full spline method (bottom) shape change is about equivalent between high (black) and low (red) altitude populations. Before correcting for allometry, we observe a posterior cross vein shift as well as a small shift in the L2 andL5 longitudinal veins. After the allometric correction, there is a shift in the position of both cross veins as well as a more prominent shift in the longitudinal L2 and L5 veins. All effects are magnified by 2x.

**Figure S2.**
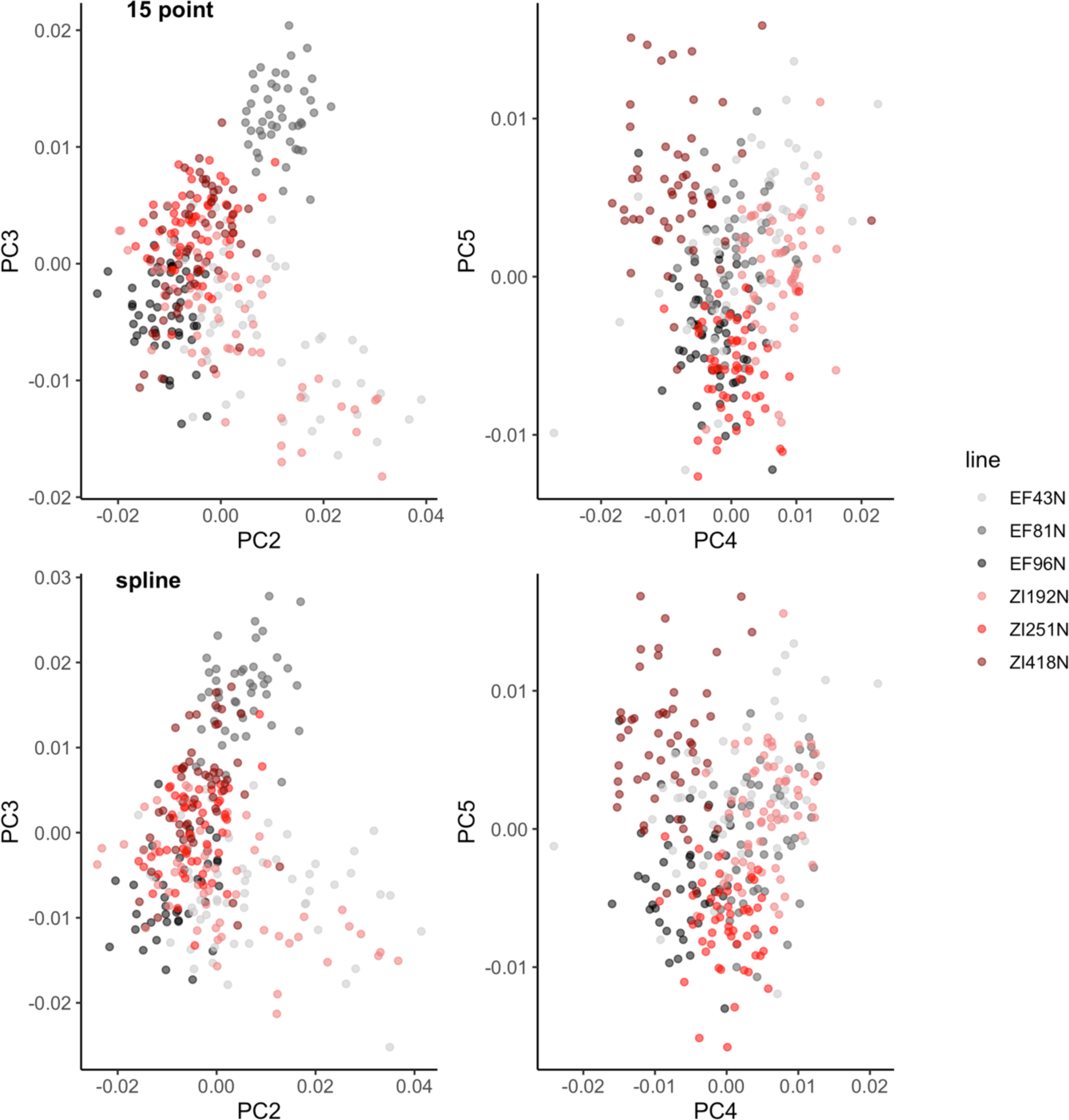
Two methods of shape collection do not substantially change the variance structure of parental populations. PCA of shape residuals for variation in high (black/greys) and low (reds) altitude populations based on inbred lines used in this analysis. PC1 is not included in this as it captures the allometric component of shape. Top panel is shape residuals captured using the 15 landmark method and bottom panel is shape residuals from the complete landmark and semi landmark method.

**Figure S3.**
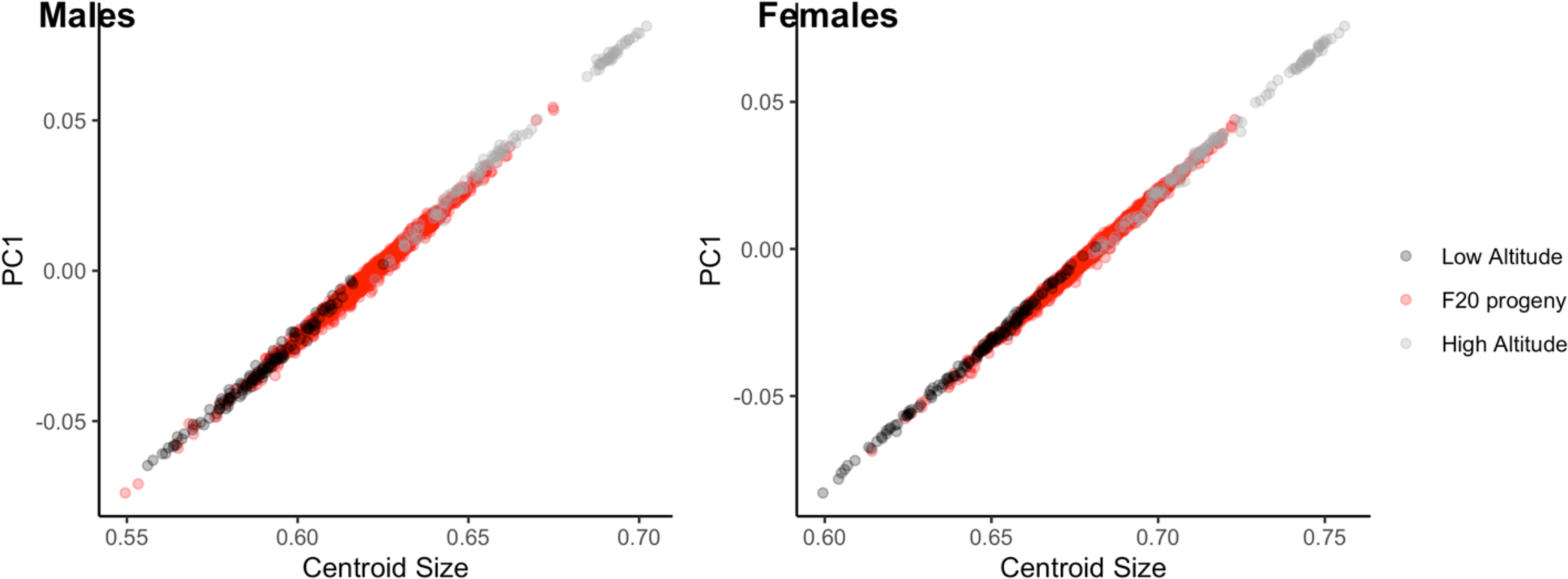
Centroid size is correlated with PC1 when log(CS) is included in PCA. Using this method, PC1 represents the allometric component of shape variation in addition to size variation. Representative data plotted for Zi192 x Ef96 cross and six inbred parental lines, but the relationship is consistent between different genotypes.

**Figure S4.**
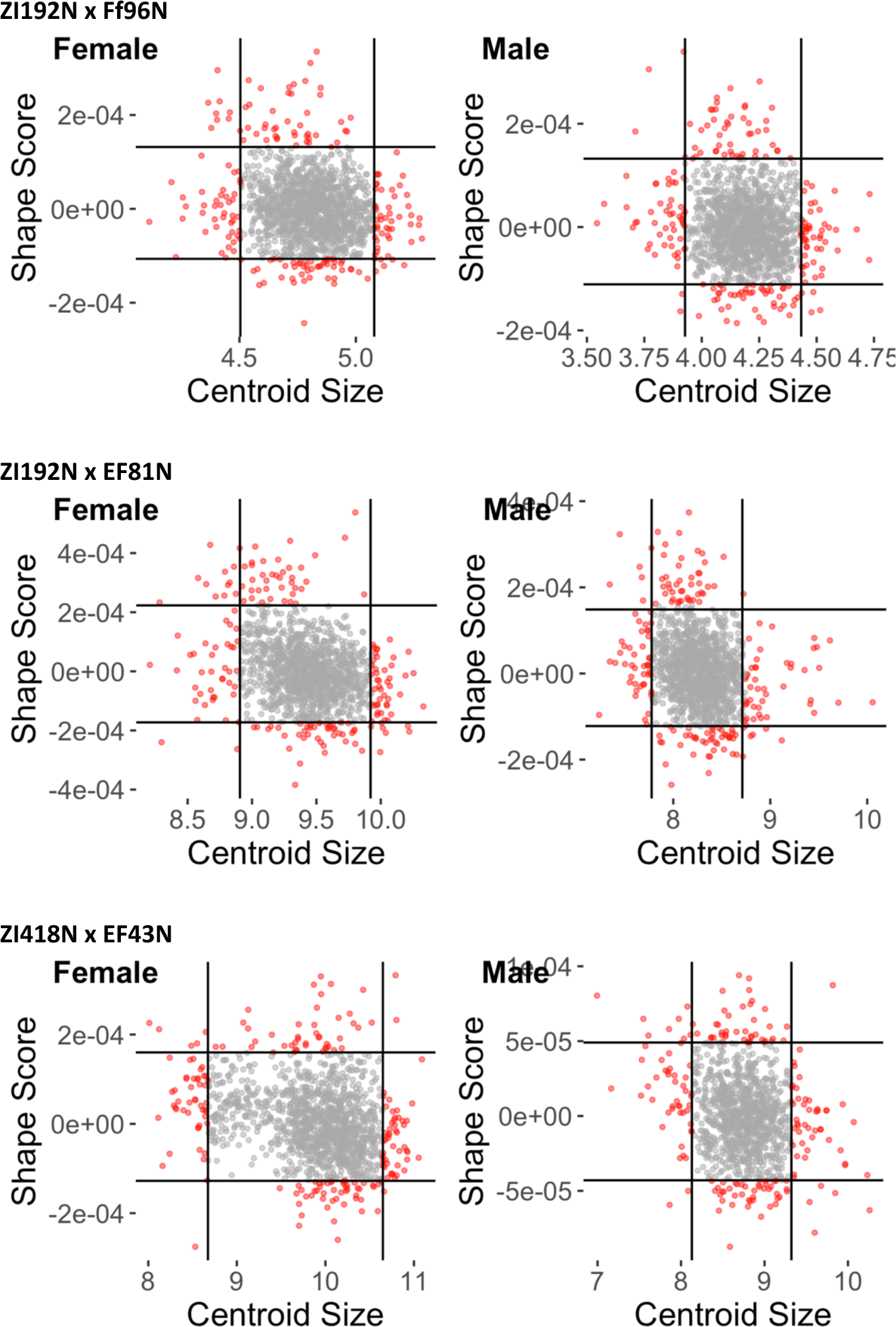
Outlier selection for creation of bulks in three F20 crosses. Each point represents an individual with those selected for sequencing indicated in red.

**Figure S5.**
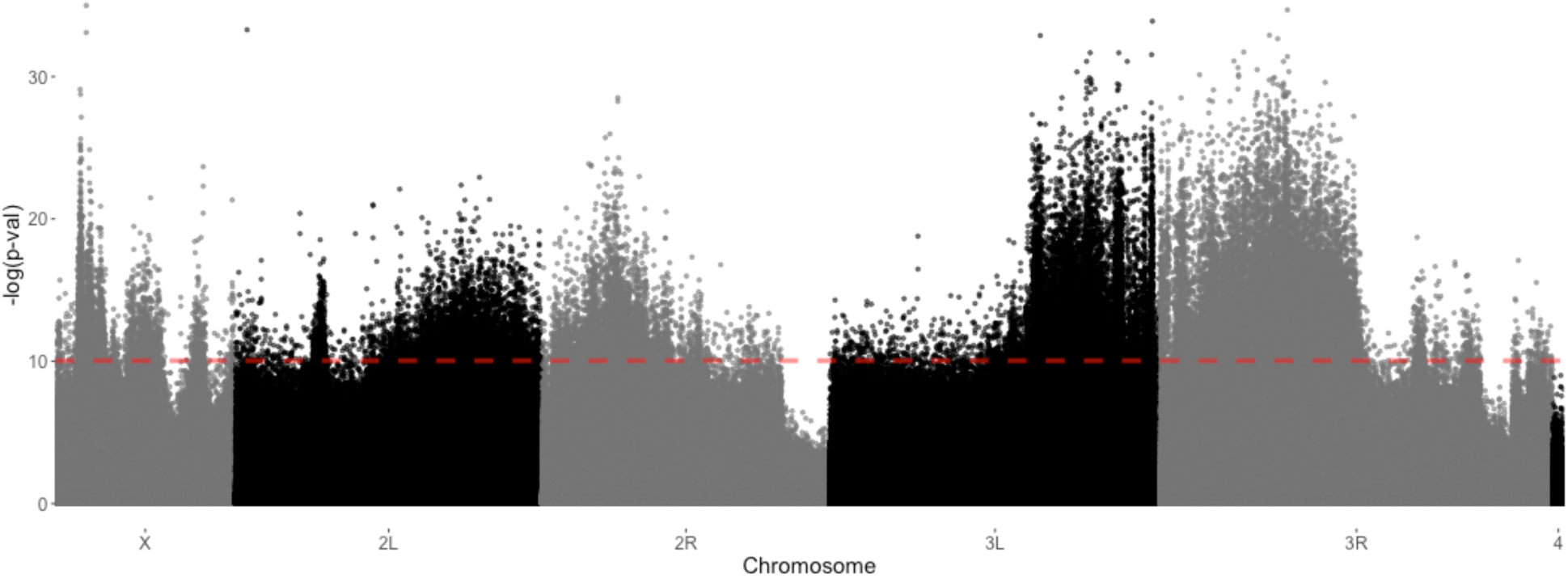
CMH test for association of SNPs with size divergence between high- and low-land populations in males. Modified CMH test to account for sampling error as implemented in the ACER (Spitzer *et al* 2020) program in R using population sizes equal to the total number of individuals phenotyped for each cross and 0 generations of selection between pools. Red line indicates a FDR of 0.01.

**Figure S6.**
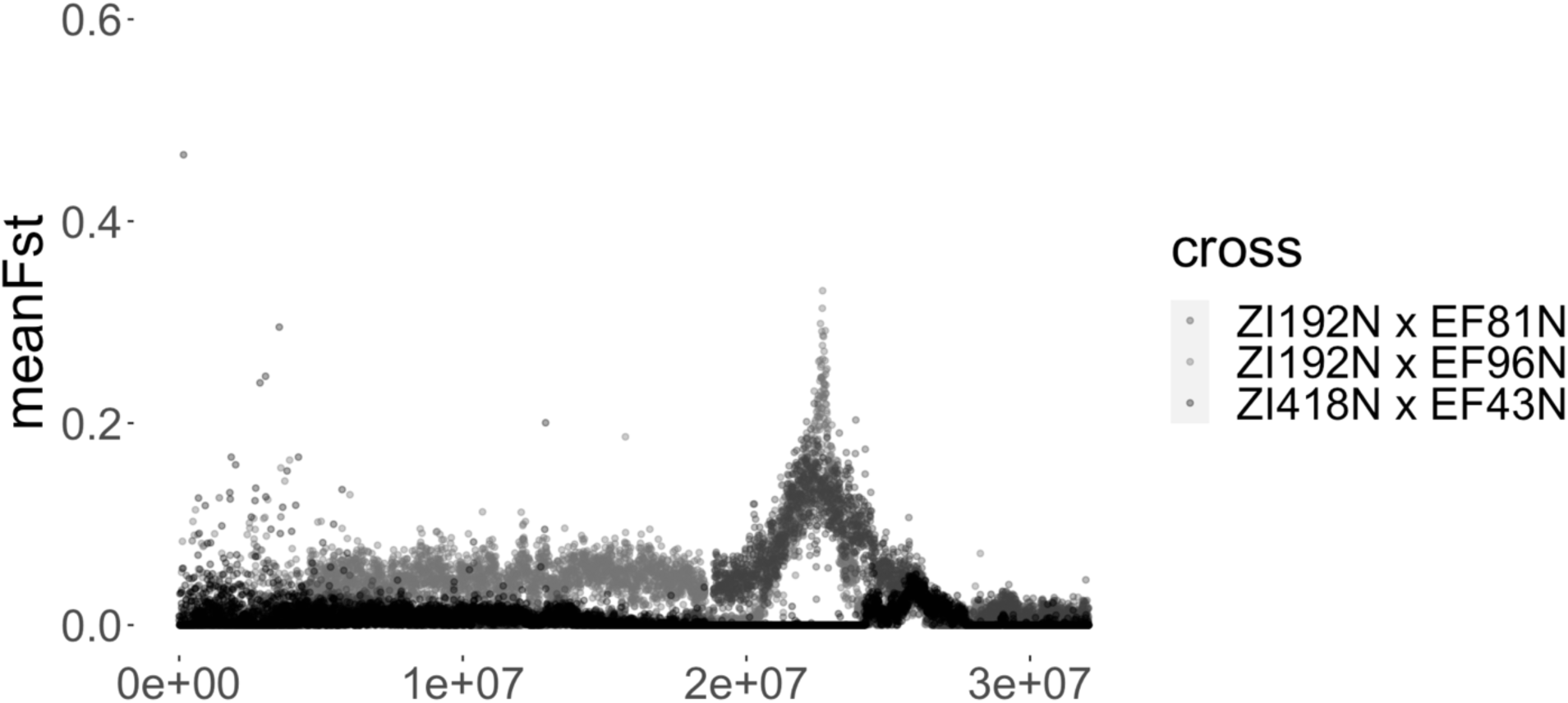
Region of genetic differentiation on chromosome 3R for shape mapping crosses. Plotting of mean F_ST_ calculated between male bulk pools in three different F20 mapping crosses, measured in 5000bp windows. X axis represents position. Shared genomic location of QTL between Zi192 x Ef81 and Zi192 x Ef96 cross can be see (greys) while small region of differentiation in Zi41 x Ef43 cross (black) is not in the same genomic region.

**Figure S7.**
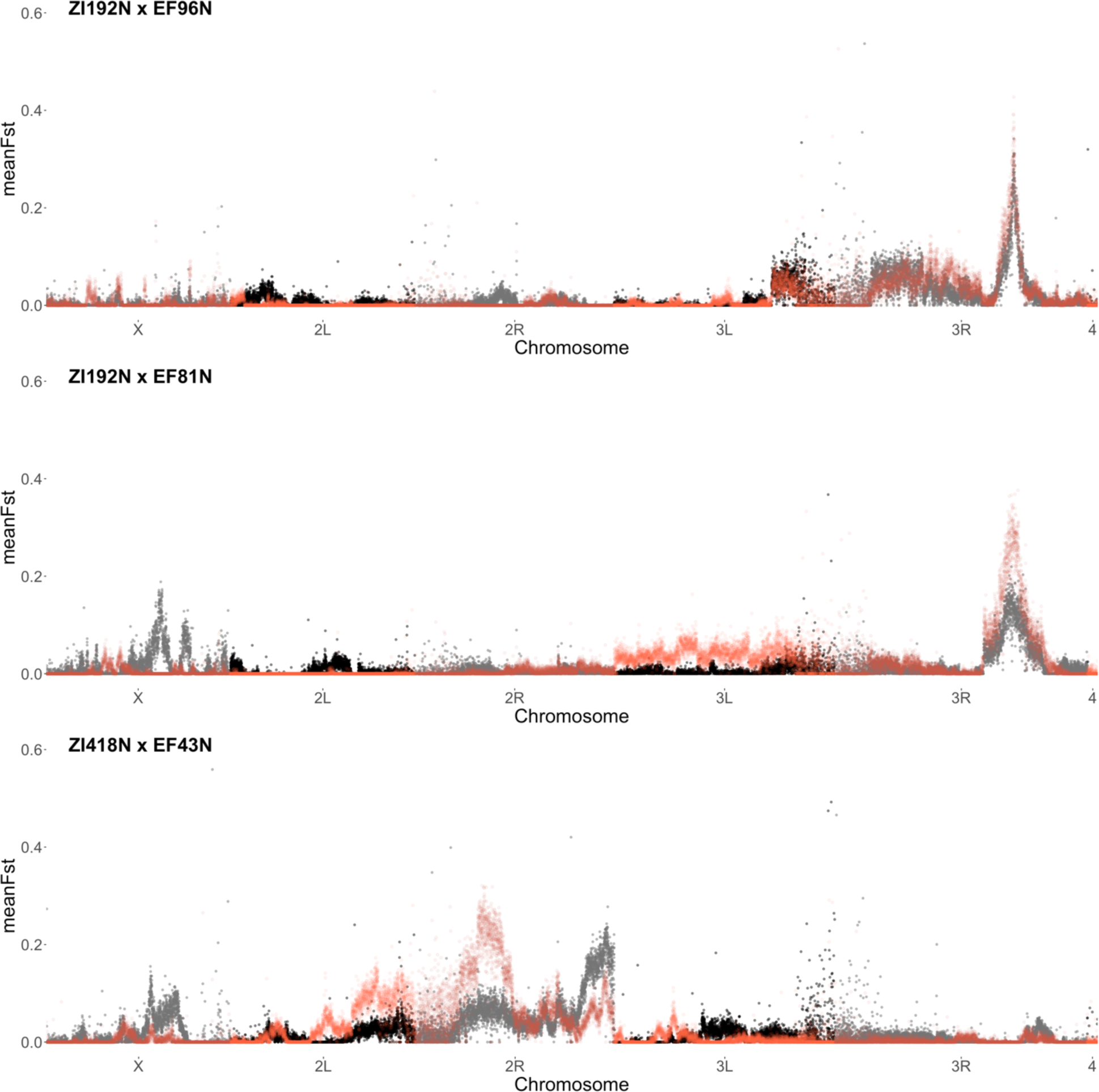
Sex specific contributions of to shape variation between high and low altitude populations. F_ST_ measured in 5000 bp windows between bulk pools within sex. Comparison between male pools (black/grey) and female pools (reds), show a similar but not completely overlapping genetic basis. This is particularly apparent on the X chromosome, which is hemizygous in males.

**Figure S8.**
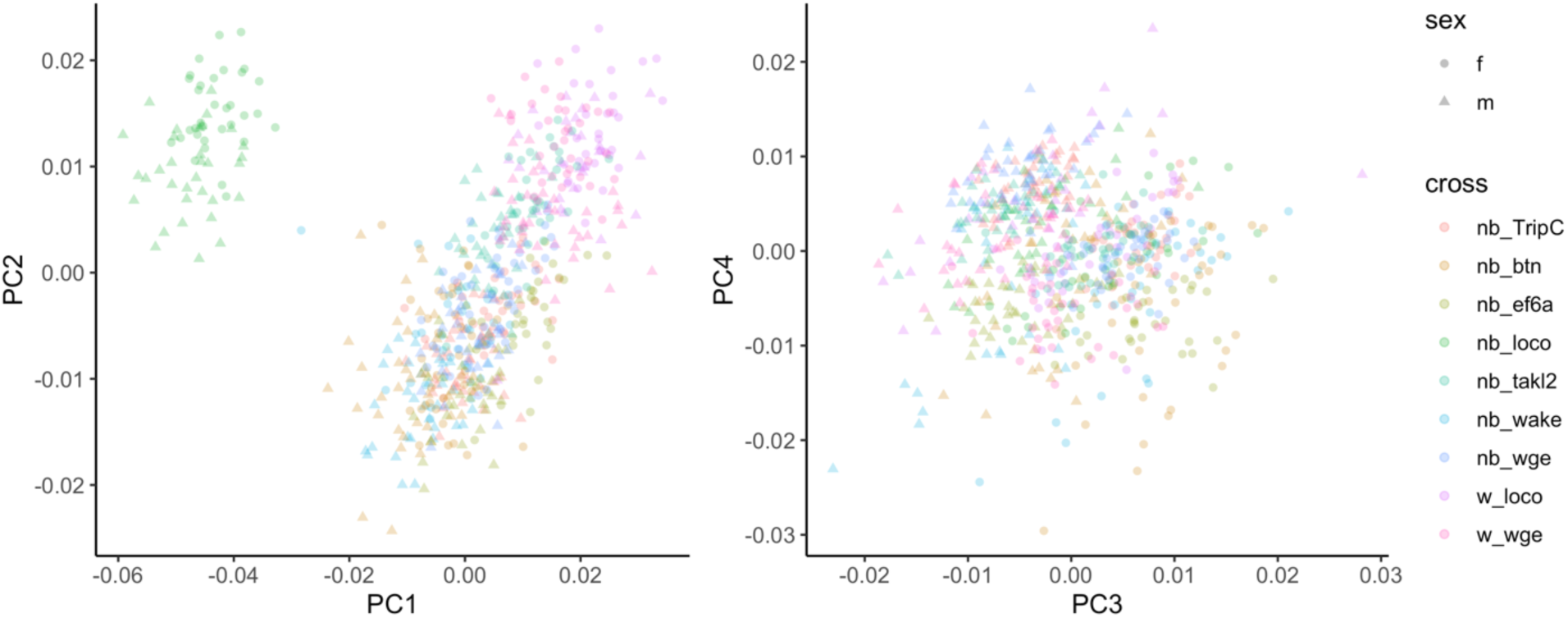
PCA of shape variation in RNAi knockdown. For the cross genotype, the *nubbin*-GAL4 line or *white*^−^ control is indicated before the underscore and the UAS-GOI-RNAi or TRiP control genotype is indicated after the underscore.

**Figure S9.**
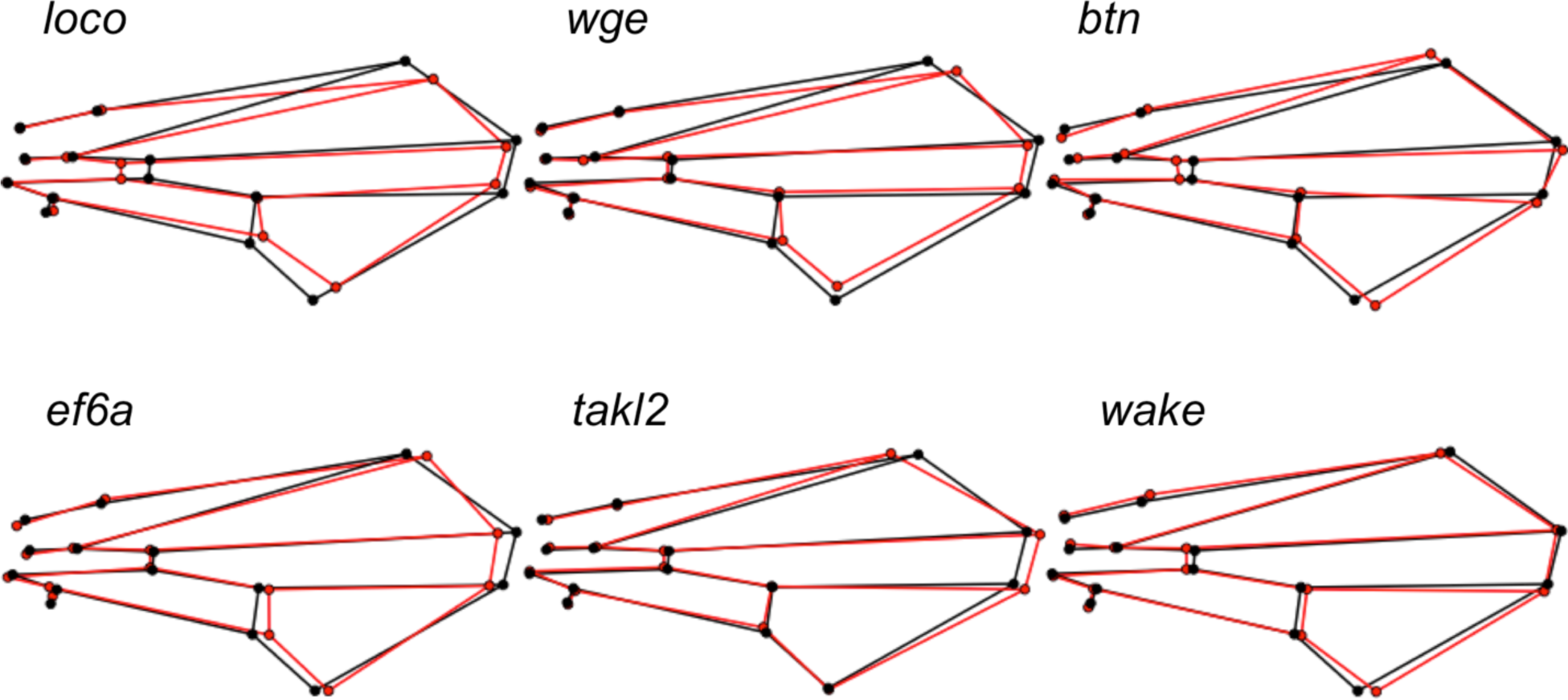
Effect on shape from knock down of candidate genes by RNAi in males. Shape change between RNAi knockdown (red) and control (black) is plotted. Effects are magnified for visualization: *loco* 1.5x; *wge* 3x; *btn* 5x; *ef6a* 2x; *takl2* 3x; *wake* 5x. Unequal magnification required due to different magnitudes of estimated effect vectors.

**Figure S10.**
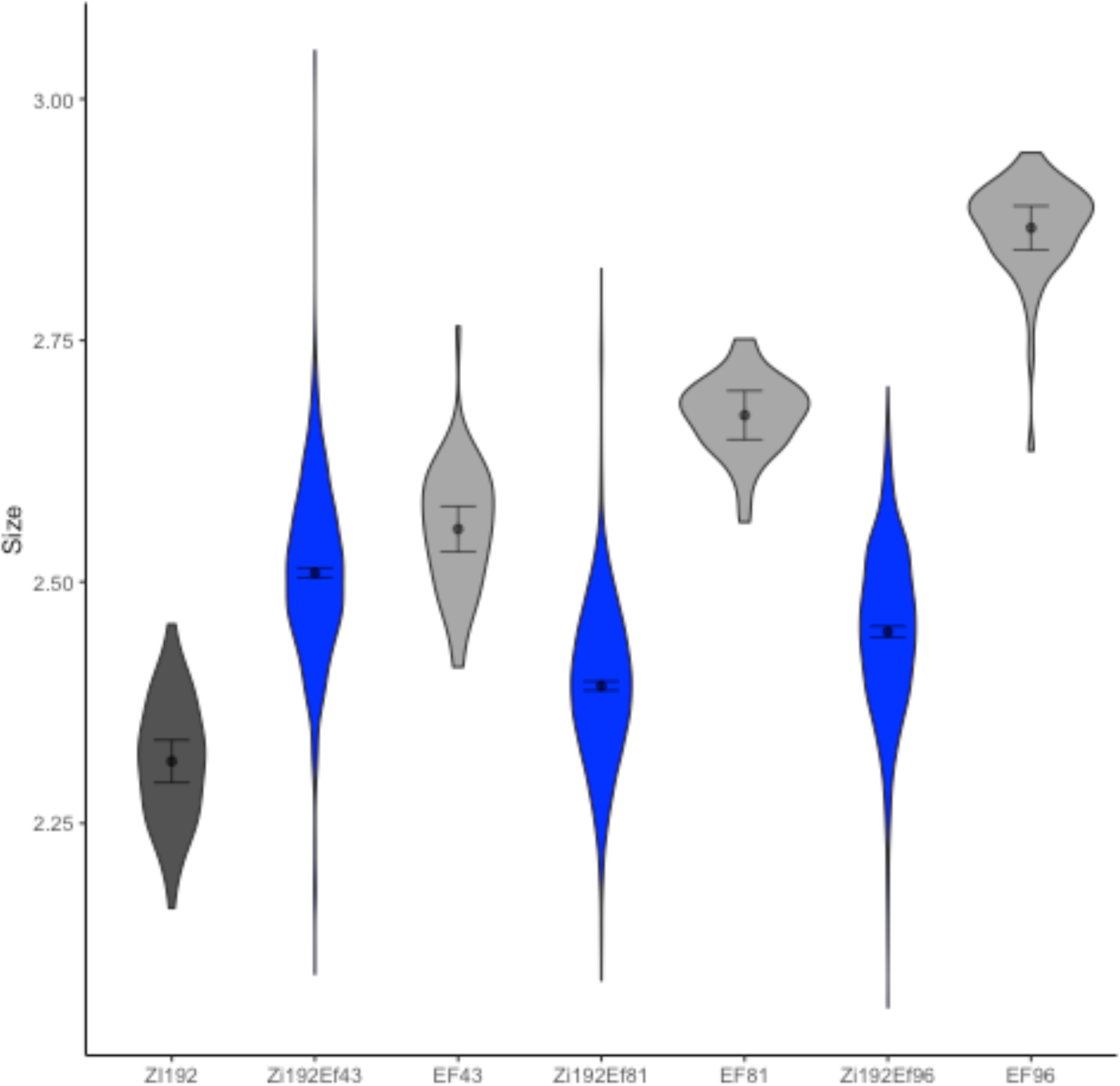
Male size variation within and between parental isogenic lines and F20 intercross males. Observed size, measured in centroid size in parental (greys) and F20 cross lines (blue). Low altitude parent is indicated in darker gray than high altitude parents. Estimated mean size with 95% CI indicated within violin plot. It should be noted that sample size is not equal between parental and cross groups, with parental lines represented by ∼50 individuals and crosses represented by >800 individuals.

**Figure S11.**
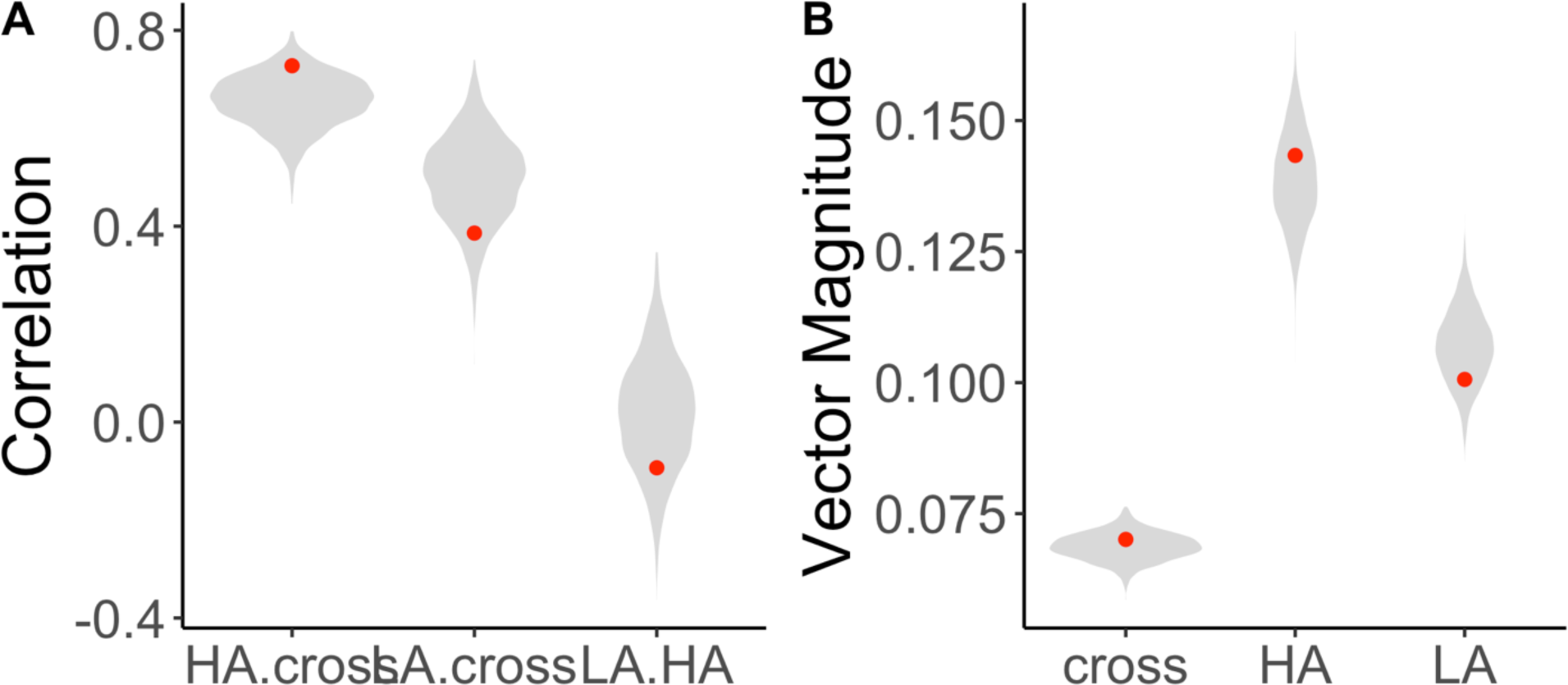
Comparison of allometry vectors between parental populations and F20 crosses. (A) Pairwise comparisons of allometric vector of shape variation between parental populations (3 HA, 3LA) or F20 cross (3 genotypes). Red points indicate observed value from data with 95% CI from 1000 bootstraps indicated in grey. (B) Magnitude (*l^2^ norm*) of allometry effect vector estimated for parental and cross genotypes. Red points indicate observed value from data with 95% CI from 1000 bootstraps indicated in grey. LA: low altitude, HA: high altitude population.

**Figure S12.**
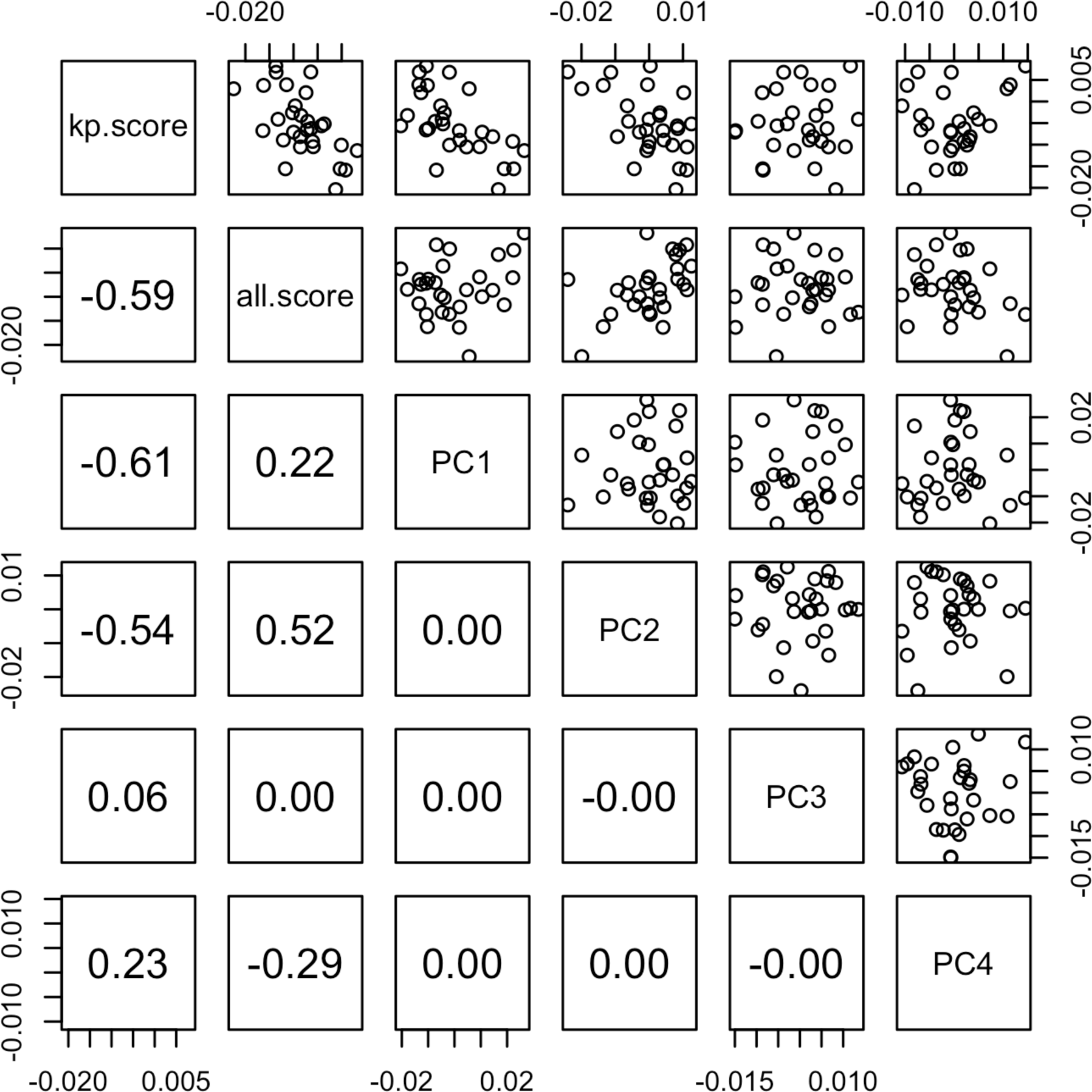
Correlation between shape scores and directions of phenotypic variation between populations in males. Principal components of shape variation estimated from allometry corrected shape residuals of landmark data from inbred African lines from this study as well as Pesevski and Dworkin 2022. Two shape scores calculated by projecting mean line shapes onto the altitude effect vector were calculated to compare the direction of shape variation assisted with adaptation to high altitude with the directions of greatest shape variation within the populations. Kp.score is calculated based only on the lines used in this paper while all.score uses all the available lines.

