## Supplementary material for "Polygenic architecture of adaptation to a high-altitude environment for *Drosophila melanogaster* wing shape and size": SuppFile1

Pool Lab *Drosophila* media:

4500 cc water
500 cc cornmeal-Quaker yellow
500 cc molasses-Grandma's unsulfured (fancy)
200 cc powdered yeast-MP Biochemical Brewers yeast
54 grams agar-Genesee Drosophila agar type II
20 ml propionic acid (100%)
45 ml tegosept 10% in 95% ethanol (4.5 g in 45 ml)

Dworkin Lab *Drosophila* media:

5.25 L water

195g Fancy Molasses (BulkBarn)

195g Blackstrap Molasses (BulkBarn)

245g Corn Meal

27g Carrageenan

50g Yeat – RedStar Baker’s Yeast

12mL propionic acid (100%)

25mL tegosept; 10% in 95% ethanol (2.5g in 25mL)
