## Supplementary material for "Polygenic architecture of adaptation to a high-altitude environment for *Drosophila melanogaster* wing shape and size": SuppFile2

Number of wings in mapping studied by genotype. The total number of wings phenotyped for each cross were: Zi192 x Ef81: Females: 1323, Males 1315; Zi192 x Ef96: Females: 1425, Males: 1432; Zi418 x Ef43: Females 1153, Males: 1262.

| Genotype | Sex | High Shape | High Size | Low Shape | Low Size | High Shape  and Size | Low Shape  and Size | High Shape  Low Size | Low Shape  High Size |
| --- | --- | --- | --- | --- | --- | --- | --- | --- | --- |
| Zi192Ef81 | Females | 43 | 45 | 39 | 37 | 5 | 11 | 2 | 0 |
|  | Males | 45 | 44 | 44 | 25 | 5 | 5 | 0 | 1 |
| Zi192Ef96 | Females | 42 | 42 | 47 | 47 | 8 | 3 | 0 | 0 |
|  | Males | 46 | 46 | 25 | 25 | 4 | 4 | 0 | 0 |
| Zi418Ef43 | Females | 48 | 42 | 48 | 49 | 1 | 6 | 1 | 2 |
|  | Males | 46 | 48 | 44 | 28 | 2 | 0 | 2 | 6 |

Number of wings used in RNAi experiment, by cross. All files were male in this analysis.

| Cross | Males | Females |
| --- | --- | --- |
| *nb*-UAS x Trip Control | 35 | 40 |
| *w^-^* x Gal4-*loco* RNAi | 28 | 33 |
| *w^-^* x Gal4-*wge* RNAi | 39 | 34 |
| *nb*-UAS x Gal4-*btn* RNAi | 36 | 38 |
| *nb*-UAS x Gal4-*ef6a* RNAi | 34 | 31 |
| *nb*-UAS x Gal4-*loco* RNAi | 33 | 33 |
| *nb*-UAS x Gal4-*takl2* RNAi | 34 | 37 |
| *nb*-UAS x Gal4-*wake* RNAi | 34 | 35 |
| *nb*-UAS x Gal4-*wge* RNAi | 37 | 37 |

Number of wings used in deletion mapping experiment, by cross. All files were male in this analysis and shape collected using 15 point method.

| Cross | Males | Deletion Pannel |
| --- | --- | --- |
| Ef119 x 25015 | 40 | DrosDel |
| Ef119 x 25693 | 19 | DrosDel |
| Ef119 x 25727 | 19 | DrosDel |
| Ef119 x 26537 | 32 | DrosDel |
| Ef119 x 27375 | 29 | DrosDel |
| Ef119 x DrosDel Control | 29 | DrosDel |
| Ef119 x 7670 | 21 | Exelis |
| Ef119 x 7671 | 20 | Exelis |
| Ef119 x 7672 | 29 | Exelis |
| Ef119 x 7740 | 18 | Exelis |
| Ef119 x Exel Control | 26 | Exelis |
| Ef43 x 25015 | 16 | DrosDel |
| Ef43x 25693 | 16 | DrosDel |
| Ef43 x 25727 | 35 | DrosDel |
| Ef43 x 26537 | 26 | DrosDel |
| Ef43 x 27375 | 0 | DrosDel |
| Ef43 x DrosDel Control | 28 | DrosDel |
| Ef43 x 7670 | 36 | Exelis |
| Ef43 x 7671 | 32 | Exelis |
| Ef43 x 7672 | 32 | Exelis |
| Ef43 x 7740 | 37 | Exelis |
| Ef43 x Exel Control | 13 | Exelis |
| Ef81 x 25015 | 28 | DrosDel |
| Ef81x 25693 | 13 | DrosDel |
| Ef81 x 25727 | 37 | DrosDel |
| Ef81 x 26537 | 20 | DrosDel |
| Ef81 x 27375 | 14 | DrosDel |
| Ef81 x DrosDel Control | 24 | DrosDel |
| Ef81 x 7670 | 15 | Exelis |
| Ef81 x 7671 | 20 | Exelis |
| Ef81 x 7672 | 30 | Exelis |
| Ef81 x 7740 | 22 | Exelis |
| Ef81 x Exel Control | 30 | Exelis |
| Ef96 x 25015 | 34 | DrosDel |
| Ef96 x 25693 | 11 | DrosDel |
| Ef96 x 25727 | 15 | DrosDel |
| Ef96 x 26537 | 25 | DrosDel |
| Ef96 x 27375 | 11 | DrosDel |
| Ef96 x DrosDel Control | 9 | DrosDel |
| Ef96 x 7670 | 0 | Exelis |
| Ef96 x 7671 | 29 | Exelis |
| Ef96 x 7672 | 30 | Exelis |
| Ef96 x 7740 | 21 | Exelis |
| Ef96 x Exel Control | 24 | Exelis |
| Zi192 x 25015 | 15 | DrosDel |
| Zi192 x 25693 | 0 | DrosDel |
| Zi192 x 25727 | 30 | DrosDel |
| Zi192 x 26537 | 27 | DrosDel |
| Zi192 x 27375 | 16 | DrosDel |
| Zi192 x DrosDel Control | 20 | DrosDel |
| Zi192 x 7670 | 28 | Exelis |
| Zi192 x 7671 | 16 | Exelis |
| Zi192 x 7672 | 14 | Exelis |
| Zi192 x 7740 | 13 | Exelis |
| Zi192 x Exel Control | 16 | Exelis |
| Zi251 x 25015 | 38 | DrosDel |
| Zi251 x 25693 | 0 | DrosDel |
| Zi251 x 25727 | 17 | DrosDel |
| Zi251 x 26537 | 29 | DrosDel |
| Zi251 x 27375 | 0 | DrosDel |
| Zi251 x DrosDel Control | 22 | DrosDel |
| Zi251 x 7670 | 14 | Exelis |
| Zi251 x 7671 | 9 | Exelis |
| Zi251 x 7672 | 19 | Exelis |
| Zi251 x 7740 | 0 | Exelis |
| Zi251 x Exel Control | 7 | Exelis |
| Zi357 x 25015 | 40 | DrosDel |
| Zi357 x 25693 | 18 | DrosDel |
| Zi357 x 25727 | 35 | DrosDel |
| Zi357 x 26537 | 29 | DrosDel |
| Zi357 x 27375 | 30 | DrosDel |
| Zi357 x DrosDel Control | 23 | DrosDel |
| Zi357x 7670 | 25 | Exelis |
| Zi357 x 7671 | 31 | Exelis |
| Zi357 x 7672 | 31 | Exelis |
| Zi357 x 7740 | 37 | Exelis |
| Zi357 x Exel Control | 34 | Exelis |
| Zi357 x 25015 | 32 | DrosDel |
| Zi357 x 25693 | 13 | DrosDel |
| Zi357 x 25727 | 31 | DrosDel |
| Zi357 x 26537 | 28 | DrosDel |
| Zi357 x 27375 | 16 | DrosDel |
| Zi357 x DrosDel Control | 34 | DrosDel |
| Zi357x 7670 | 23 | Exelis |
| Zi357 x 7671 | 26 | Exelis |
| Zi357 x 7672 | 34 | Exelis |
| Zi357 x 7740 | 25 | Exelis |
| Zi357 x Exel Control | 18 | Exelis |

Number of wings used in shape variation analysis, all wing shape was collected using the 15 point method. Only males were used for this analysis. HA = high altitude, LA = low altitude.

| Genotype | Males | Group |
| --- | --- | --- |
| EF43 | 53 | HA |
| EF81 | 45 | HA |
| EF96 | 56 | HA |
| ZI192 | 60 | LA |
| ZI251 | 52 | LA |
| ZI418 | 53 | LA |
| Zi192 x Ef43 | 1185 | Cross |
| Zi192 x Ef81 | 1288 | Cross |
| Zi192 x Ef96 | 880 | Cross |

Number of wings used in shape used to estimate direction of altitudinal effect shape change. Data collected by both Maria Pesevski(from Pesveski and Dworkin 2022) and as part of this work. Only data for Zi251 was collected in both studies, as indicated with an *. Lines from this study are bolded. All data was collected using the WingMachine pipeline. HA = high altitude, LA = low altitude.

| Population | Line | Males | Females |
| --- | --- | --- | --- |
| HA | Ef122 | 14 | 11 |
| HA | EF131 | 35 | 22 |
| HA | Ef134 | 16 | 18 |
| HA | Ef136 | 20 | 17 |
| HA | Ef15 | 24 | 19 |
| HA | Ef19 | 13 | 10 |
| HA | Ef39 | 21 | 11 |
| HA | **Ef43** | 40 | 48 |
| HA | Ef65 | 13 | 19 |
| HA | Ef73 | 26 | 17 |
| HA | **Ef81** | 32 | 39 |
| HA | **Ef96** | 33 | 44 |
| HA | Ef16 | 25 | 26 |
| HA | Ef217 | 31 | 33 |
| HA | Ef98 | 20 | 15 |
| LA | Zi124 | 21 | 18 |
| LA | Zi159 | 29 | 34 |
| LA | Zi160 | 19 | 22 |
| LA | **Zi192** | 46 | 49 |
| LA | Zi216 | 28 | 19 |
| LA | Zi217 | 14 | 12 |
| LA | **Zi251*** | 68 | 60 |
| LA | Zi254 | 47 | 38 |
| LA | Zi311 | 16 | 21 |
| LA | Zi367 | 22 | 15 |
| LA | **Zi418** | 52 | 50 |
| LA | Zi455 | 22 | 12 |
| LA | Zi461 | 26 | 39 |
